# A hippocampal neuroimaging signature of neurovascular insulin signalling links metabolism to mood

**DOI:** 10.64898/2026.06.17.732833

**Authors:** Antoine Cherix, Beata R Godlewska, Alberto Lazari, Sijia Zhao, Gabrielle Dugan, Mohamed Tachrount, Maria Greco, Sean Smart, William T Clarke, David Bannerman, Charlotte J Stagg, Masud Husain, Phil Cowen, Jason P Lerch

## Abstract

Depression is a leading cause of global disability and is increasingly linked to systemic metabolic dysfunctions, including insulin resistance. However, the biological pathways connecting metabolic state to affective symptoms are unresolved. A growing body of evidence indicates that insulin, beyond its role in systemic glucose homeostasis, supports brain metabolism, synaptic function and cognition, yet its contribution to mood regulation remains unclear.

Here, we identify a human hippocampal metabolic signature associated with glycaemic variation in individuals with depression. Using multimodal neuroimaging, we show that hippocampal concentrations of GABA and lactate correlate with HbA1c and with mood severity, while functional connectivity between the hippocampus and the default mode network tracks affective symptoms independently of adiposity. To causally probe these relationships, we generated a mouse model of hippocampus-specific insulin receptor depletion. Unexpectedly, reducing insulin signalling at the blood brain barrier enhanced neuronal metabolism and attenuated anxiety-like behaviour. Notably, hippocampal lactate and GABA levels similarly tracked anxiety-related behaviour in mice, mirroring their association with symptom severity in humans.

Together, these findings identify a conserved neurometabolic signature linking hippocampal insulin signalling to affective state, and reveal that brain insulin resistance exerts context- and cell-type-specific effects on behaviour. This work establishes a mechanistic basis for brain insulin resistance in depression and highlights the hippocampus as a critical hub for metabolic modulation of mood.

## Introduction

Depression is among the leading causes of disability worldwide and represents a major public health challenge^1,2^. Increasing evidence links depression to metabolic dysfunction, including insulin resistance^3–8^, yet the biological pathways connecting these conditions remain unresolved. Insulin is a key anabolic hormone that regulates systemic glucose homeostasis, and emerging evidence indicates that insulin signalling in the brain supports metabolism, synaptic plasticity^9–11^ and cognition^12,13^. These observations have led to the concept of *brain insulin resistance* (BIR)^14,15^, broadly defined as a reduced responsiveness of the central nervous system to insulin, and proposed as a mechanistic link between metabolic and neuropsychiatric disorders^16^.

Support for this model derives largely from rodent studies in which the insulin receptor is deleted globally or in specific neural cell types^17^. These models demonstrate that insulin signalling is required for metabolic regulation and plasticity within the brain, and highlight the importance of tightly controlled insulin access and action across the blood-brain barrier (BBB). However, translating these findings to humans has been challenging, in part due to the lack of non-invasive approaches to assess insulin signalling in the living brain^18^. As a result, it is unknown whether brain insulin resistance represents a uniform deficit in neuronal insulin action, or a more complex, region- and cell-type-specific process. Consequently, the mechanisms through which brain insulin resistance influences affective function and contributes to depression remain poorly understood.

We previously identified a neuroimaging signature in a mouse model of comorbid depression and metabolic syndrome that implicated the hippocampus as particularly vulnerable to hyperglycaemia and hyperinsulinaemia^19^. This phenotype was characterised by depressive-like behaviour, neuroenergetic impairments, and alterations in excitatory and inhibitory metabolism measured with magnetic resonance spectroscopy (MRS).

Here, we test whether this metabolic signature is present in individuals with depression and whether it relates to systemic glycaemic control. We further examine its mechanistic basis using a mouse model of localised insulin receptor depletion in the hippocampus. We show that hippocampal metabolism and functional connectivity track glycaemic variation and mood symptoms in humans. Unexpectedly, selective reduction of insulin signalling in non-parenchymal (vascular) hippocampal cells enhances neuronal metabolic function and attenuates anxiety-related behaviours in mice. Together these findings identify a hippocampal neuroimaging signature of altered insulin signalling and reveal that BIR can operate as a cell-type-specific, context-dependent process linking metabolism to mood.

## Results

### Fingerprinting hippocampal response to glycaemia in depression

To investigate whether the hippocampus is sensitive to metabolic variation associated with insulin resistance, we first examined hippocampal integrity in individuals with depression using multimodal neuroimaging on 7 Tesla MRI scanner (anatomical MRI, MRS and resting-state fMRI), and related these measures to metabolic indices, mood-related symptoms and cognition (Fig. 1). The study included nine currently-unmedicated individuals meeting criteria for major depressive disorder and twelve healthy controls which were comparable in age and sex distribution (Supplementary Table 3). No participant had a history of diabetes or major metabolic disorders. Compared with healthy controls, the depressed group exhibited significantly higher hippocampal concentrations of GABA and lactate (Fig.1a-e). Notably, participants with depression also showed elevated – though clinically normal – levels of HbA1c, a marker of sustained glycaemia and systemic insulin resistance Fig.f). In contrast, no group differences were observed in metabolite concentrations within the dorsal anterior cingulate cortex (dACC; Supplementary Fig.S1a-e), a region often implicated in depression^20^, or in hippocampal volume (Supplementary Fig.S1f). The depressed group showed significantly higher functional connectivity between the hippocampus and the default mode network (DMN), with a stronger effect observed in the anterior compared to the posterior hippocampus (Fig.1g). Despite robust hippocampal neurochemical differences, groups did not differ in BMI, % body fat, or visceral fat (Supplementary table 3), indicating that the observed brain metabolic signature is dissociable from adiposity.

**Fig. 1:**
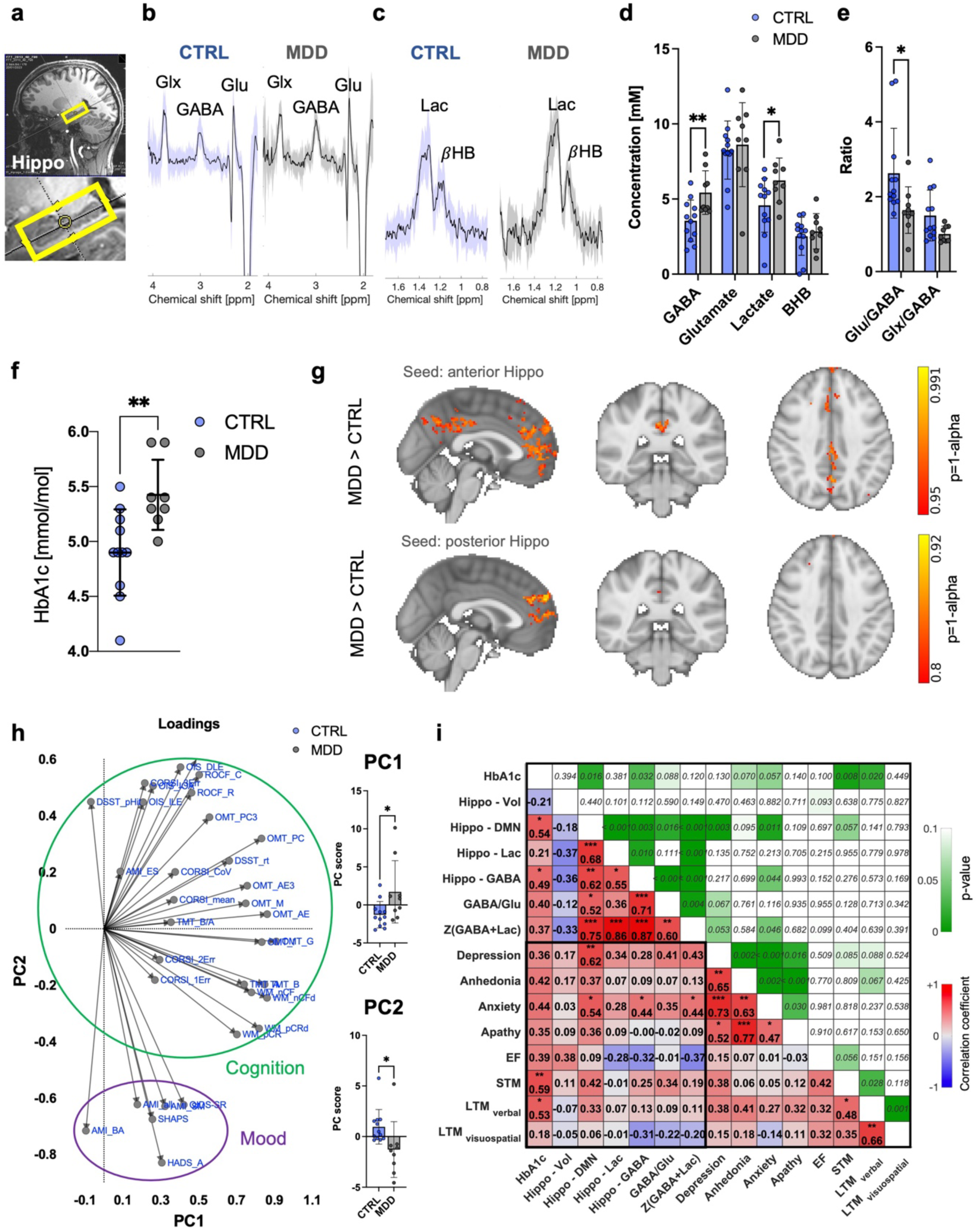
Hippocampal function and metabolism are associated with glycaemic variation and mood symptoms in depression. **a.-c.** Hippocampal (**a**) voxel placement and average spectra (±s.d) for (**b**) GABA-edited and (**c**) lactate-edited ^1^H-MRS in CTRL (N=12, blue) and MDD (N=9, grey) groups. **d.-e.** Quantification of hippocampal metabolites from ^1^H-MRS spectra. Student’s t-test, *p<0.05, **p<0.005. **f.** Group differences in blood HbA1c values. Student’s t-test, *p<0.05, **p<0.005. **g.** Anterior (top) and posterior (bottom) hippocampal resting-state functional connectivity. **h.** Variable PCA loadings showing the separation between cognitive performance metrics and mood-related measures. Participant scores illustrating group differences between individuals with depression and healthy controls along the derived behavioural dimensions. Principal component 1 (PC1) and 2 (PC2) explained 31.4% and 17.4% of the variance respectively. Student’s t-test, *p<0.05 **i.** Exploratory analysis of key brain functional metrics correlation with cognition and mood. Correlation coefficient is given as Spearman’s r and p-values were not corrected for multiple comparisons. LTM, long -term memory; STM, short-term memory; EF, executive functions; Lac, lactate; DMN, default-mode network; Vol, volume.

Principal component analysis of 34 standardized behavioural variables identified two components explaining 48.8% of the total variance (PC1: 31.4%, PC2: 17.4%). Loadings revealed a clear dissociation between cognitive and mood measures, with cognitive performance loading predominantly onto positive PC1/PC2 values and mood-related variables loading negatively on PC2, indicating a largely separable affective dimension (Fig.1h). Component scores differed between individuals with depression and healthy controls (PC1: MDD > CTRL, p=0.033; PC2: CTRL > MDD, p=0.032). Neuroimaging markers showed stronger association (Fig.1i) with affective symptoms (defined here as depression and anxiety severity) than with cognitive measures. Relationships were most pronounced for depression severity (FC_hippo-DMN_ vs. QIDS-SR: r=.62; Z_GABA+Lac_ vs. QIDS-SR r=.43) and anxiety severity (FC_hippo-DMN_-HADS-A: r=.54; Z_GABA+Lac_ vs. HADS-A: r=.44), whereas associations with cognitive performance were generally weaker and non-significant. This included short-term memory and long-term verbal memory, which showed the strongest relationships with affective symptoms (e.g. STM vs. QIDS-SR: r=.38; LTM_verbal_ vs. QIDS-SR: r=.38). HbA1c was associated with most mood-related symptoms (r=.35-.45), cognitive performance (r=.39-.59), and neuroimaging measures (r=.37-.54), suggesting a broad contribution to behavioural and brain phenotypes. Together, these findings indicate that depression-related variability is structured along partially dissociable affective and cognitive axes, with neuroimaging markers of hippocampal metabolism and function preferentially tracking affective symptom severity.

### A mouse model of hippocampal insulin resistance

To test how impaired hippocampal insulin signalling at the neurovascular interface affects metabolism and the associated neuroimaging signature, we generated an inducible model of localized brain insulin resistance (iNIRKO), in which *Insr* deletion is driven by Nestin expression^21^ in adulthood (Cre-loxP recombination; Fig.2a). Given the uncertain cellular specificity of Nestin-driven recombination with this Nes-CreERT2 strain (see Methods), we quantified *Insr* loss using both bulk (RT-qPCR) and cell-type-specific (snRNA-seq) approaches.

**Fig 2:**
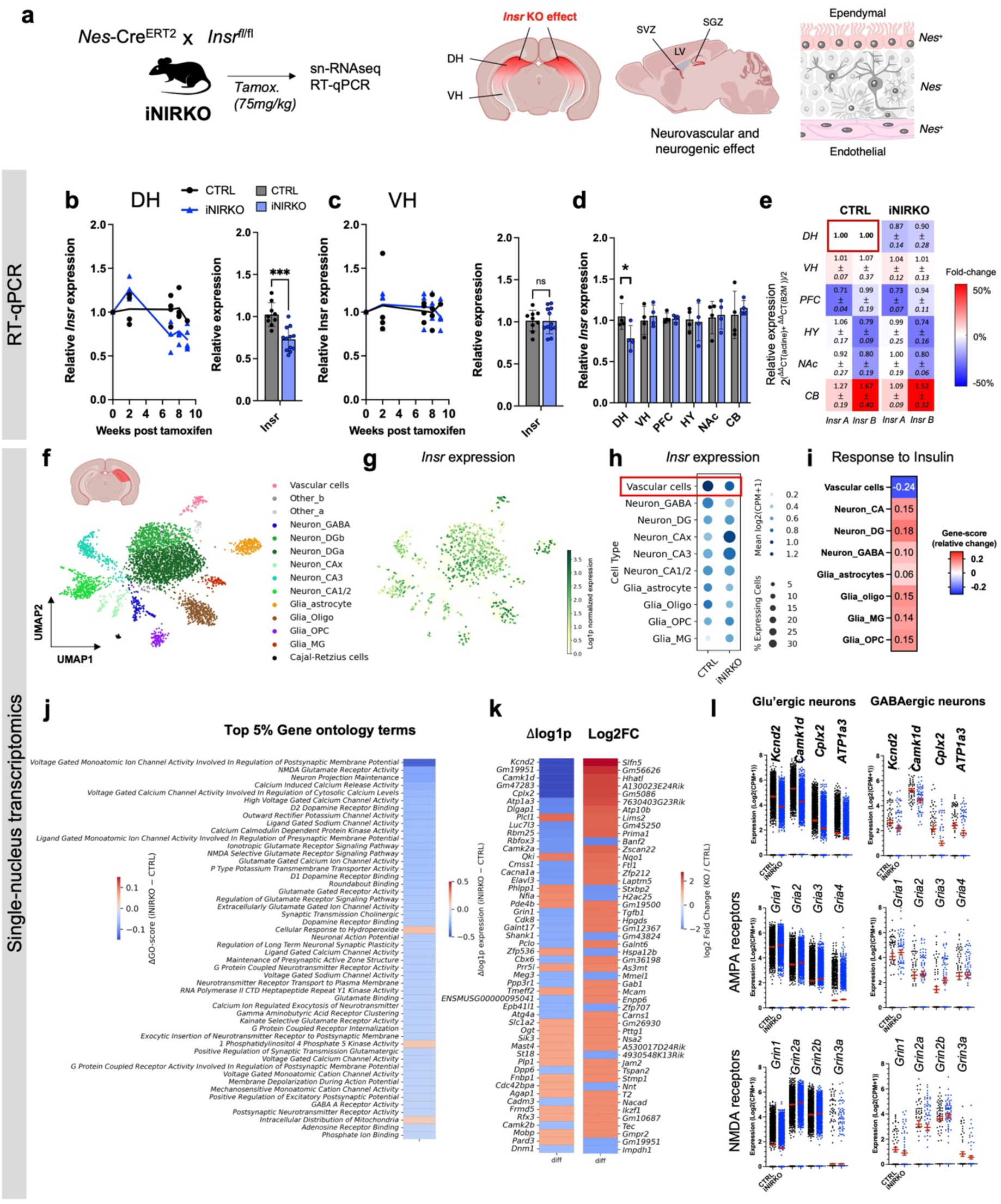
Non-parenchymal insulin receptor loss in the hippocampus models local brain insulin resistance and enhances neuronal insulin-responsive gene expression. **a.** iNIRKO mice were generated using a tamoxifen-inducible Cre line under Nestin promoter bred with floxed insulin receptor mice. Gene expression was monitored using single nucleus RNAseq and quantitative PCR (RT-qPCR). **b, c.** Timecourse of Insr expression in the dorsal (DH) and ventral (VH) parts of the hippocampus of iNIRKO mice following tamoxifen administration, measured with qPCR. iNIRKO, n=12; CTRL, n=9. **d.** Region-specificity of Insr expression at 8 weeks post tamoxifen administration. n=4 animals per group and 4 Insr primers averaged per animal. **e.** Region-specific expression of Insr isoform A and B relative to dorsal hippocampus (DH) expression **f.,g.** Uniform manifold approximation and projection (UMAP) of single nucleus RNAseq of dorsal hippocampus with (**g**) Insr expression for CTRL (n=1,674 cells, 5 pooled samples) and iNIRKO (n=2,258 cells, 6 pooled samples) mice. **h.** Dotplot of Insr gene expression across cell types in the dorsal hippocampus. **i.** Relative change (iNIRKO-CTRL) in gene score for the reponse to insulin GO term. **j.** Top 50 (5%) GO terms with highest difference in gene score (iNIRKO-CTRL) ordered by absolute difference magnitude **k. .** Top 50 genes with highest change in gene expression (iNIRKO-CTRL) reported as difference (Δlog1p) or fold change (Log2FC) **l.** Average expression of selected genes for either excitatory or inhibitory neurons. Data are mean ± s.e.m DH, dorsal hippocampus; VH, ventral hippocampus; PFC, prefrontal cortex; HY, hypothalamus; NAc, nucleus accumbens; CB, cerebellum; MG, microglia; OPC, Oligodendrocyte precursor cells; Subfields of the hippocampus: CA, Cornu ammonis; and DG, Dentate gyrus. **b.,c.,d.,** Data are mean ± s.d. Unpaired Student’s t-test. *p <0.05; ***p <0.0005; ns, not significant.

Six weeks after tamoxifen induction, iNIRKO mice showed a 20-30% reduction in bulk *Insr* expression selectively in the dorsal hippocampus, with no changes detected in other brain regions, affecting both receptor isoforms (Fig.2b-e). Single-nucleus transcriptomics identified vascular cells as the primary contributors to this reduction (Fig.2f-h; Supplementary Fig.S2), with endothelial populations – which typically express high levels of *Nes* and *Insr* – showing the strongest decrease. This was corroborated by reduced activity of insulin-responsive expression pathways (GO: response to insulin) specifically in vascular cells (Fig.2i).

Unexpectedly, reduced vascular *Insr* expression was accompanied by increased *Insr* mRNA levels and enhanced insulin-responsive transcription in parenchymal cells, most prominently neurons (Fig.2i).

Gene ontology and transcript-level analyses revealed coordinated changes in neurotransmission-related pathways across cell types, including reduced expression of *Kcnd2, Camk1d, Cplx2* and *ATP1a3* (Fig.2j-l). In parallel, *Glut1* expression was decreased in vascular cells (Supplementary Fig.S2m).

Together these findings are consistent with a compensatory increase in neuronal insulin sensitivity following reduced insulin signalling at the blood-brain-barrier.

### Hippocampal insulin resistance enhances metabolism and reduces anxiety-like behaviour

To determine the metabolic, functional, and behavioural consequences of the partial loss of insulin signalling at the hippocampal vasculature in iNIRKO mice, we performed behavioural testing followed by *in vivo* MRI and MRS 6 weeks after tamoxifen-induced recombination (Fig.3a). Behaviour was tested using established exploratory conflict-based tests of anxiety, including the elevated plus maze (EPM), light-dark box (LDB), and aversive open-field (AOF) tests, which rely on the animal’s natural aversion to open or brightly lit spaces. iNIRKO mice displayed a robust reduction in trait anxiety across multiple paradigms, including the EPM, LDB and AOF, an effect observed in both sexes (Fig. 3b–c). Consistent with this anxiolytic phenotype, iNIRKO mice showed reduced immobility and faecal output during the tests, without alterations in locomotor activity or rearing behaviour as tested with an independent locomotor activity (LMA) test (Supplementary Fig. S3a-e).

**Fig. 3.**
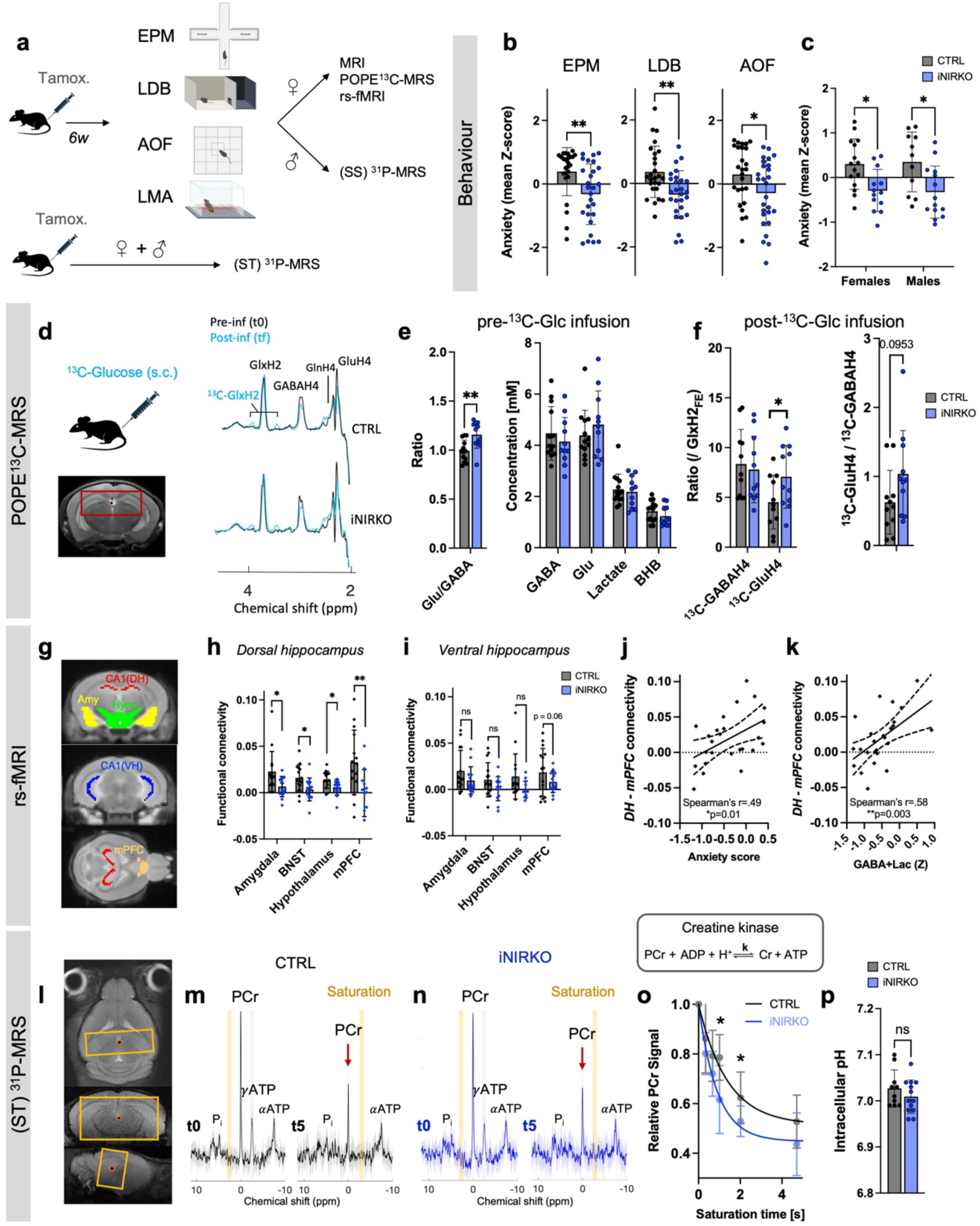
Non-parenchymal insulin resistance increases metabolic activity and function in the hippocampus and has anxiolytic-like effects. **a.** Schematic of behavioural and neuroimaging protocol. **b.,c.** Anxiety z-score in (**b**) individual EPM, LDB and AOF tests or (**c**) averaged across all tests. (**b**) Student’s t-test, *p<0.05, **p<0.005; (**c**) Two-way ANOVA, Genotype effect: F1,49 = 16.03, P = 0.0002; followed by Bonferroni’s test, *p < 0.05. **d.** POPE^13^C-MRS hippocampal voxel and averaged spectra before (black) and after (blue) a 1h20min infusion of ^13^C-glucose (U-^13^C_6_]Glc). Structural MRI with voxel placement (red) for the POPE^13^C-MRS acquisition. **e.,f.** Quantification of hippocampal metabolites (**e**) before and relative labelling (**f**) after ^13^C-glucose infusion. No labelling in the lactate or BHB pools were detected. Labelling ratios of GABAH4 and GluH4 are reported relative to the fractional enrichment (FE) of GlxH2 (left) or relative to each-other (right). Student’s t-test, *p<0.05, **p<0.005. **g.,h.,i.,** Resting state functional MRI (rs-fMRI) connectivity (Pearson’s correlation coefficient) of the hippocampus with key anxiety-related brain structures. Student’s t-test after r-to-z conversion, *p<0.05, **p<0.005, ns: not significant. **j.,k,** Relationship of dorsal hippocampus-to-mPFC connectivity with anxiety score (correlation, *p<0.05) and with metabolite concentration reported as averaged z-score of GABA and lactate concentrations (correlation, **p<0.005). Simple linear regression fit shown with 95% CI. **l.-p.** Saturation-transfer (ST) ^31^P-MRS experiments with (**l**) voxel placement and averaged spectrum (± s.d) for (**m**) CTRL and (**n**) iNIRKO mice at minimum (t0) and maximum (t5) saturation times. Fit of relative PCr signal with increasing saturation time to extract the creatine kinase forward rate constant (k). **p.** Intracellular pH measured from chemical P_i_-PCr chemical shift difference measured in steady-state (SS) ^31^P-MRS. Tamox, Tamoxifen; EPM, Elevated plus maze; LDB, light-dark box; AOF, Aversive open field; LMA, Locomotor activity; MRS, magnetic resonance spectroscopy; rs-fMRI, resting-state functional MRI; (SS) ^31^P-MRS, steady-state ^31^P-MRS; (ST) ^31^P-MRS, saturation transfer ^31^P-MRS; GABA, gamma-aminobutyric acid; Glu, glutamate; BHB, beta-hydroxybutyrate. Amy, Amygdala; Hypo, hypothalamus; mPFC, medial prefrontal cortex; CA1(DH), Cornu ammonis 1 part of the dorsal hippocampus; CA1(VH), Cornu ammonis 1 part of the ventral hippocampus; BNST, Bed nucleus of the stria terminalis; P_i_, inorganic phosphate; PCr, phosphocreatine; (α, β and γ)-ATP, adenosine triphosphate; ADP, adenosine triphosphate; Cr, creatine.

To assess hippocampal metabolism, we used POPE^13^C-MRS, which non-invasively quantifies neurotransmitter-specific glucose metabolism, as well as basal tissue content of key metabolites (glutamate, GABA, lactate, and β-hydroxybutyrate)^22^. POPE^13^C-MRS tracks metabolite’s relative ^13^C enrichment following ^13^C-glucose administration, providing an index of glucose uptake and utilisation across excitatory and inhibitory neuronal pools^22^ (Fig.3d). iNIRKO mice exhibited increased ^13^C labelling of glutamate relative to GABA, indicating a shift toward enhanced glutamatergic and/or mitochondrial metabolism (Fig.3e-f). In parallel, resting-state fMRI revealed a marked reduction in functional connectivity (Fig.3g-i) between the hippocampus and key anxiety-related regions, including the amygdala, bed nucleus of the stria terminalis (BNST), hypothalamus, and medial prefrontal cortex (mPFC). This effect was most pronounced for dorsal hippocampus–mPFC connectivity, which correlated positively with anxiety-related behaviour, reinforcing the role of hippocampal–prefrontal coupling in anxiety regulation (Fig.3j-k). A similar, albeit non-significant, trend was observed in the ventral hippocampus (Fig.3i).

To examine the neuroenergetic consequences of vascular *Insr* depletion, we used ^31^P-MRS to quantify high-energy phosphate metabolites and creatine kinase (CK) flux, a measure of mitochondrial ATP turnover (Fig.3l-n). While concentrations of ATP, inorganic phosphate (Pi), and phosphocreatine (PCr) were comparable between groups (Supplementary Fig.S3f-g), the CK forward rate constant was significantly increased in iNIRKO mice (95% CI: 0.25-46 s^-1^ (CTRL) vs. 0.50-74 s^-1^ (iNIRKO)), indicating elevated hippocampal ATP turnover (Fig.3o). This increase was not driven by alterations in intracellular pH, which can modulate CK flux (Fig.3p). Unexpectedly, total NAD(H) levels (NAD_tot_) were reduced in iNIRKO mice and correlated with anxiety-related behaviour (Supplementary Fig.S3f-i). While genes involved in NAD biosynthesis (*Nampt*, *Nmnat1*) were upregulated, gene ontology analysis instead suggested enrichments of pathways related to nucleosidase activity (Supplementary Fig.S3j-k). This signal was associated with genes linked to interleukin-1 signalling, including *Il1rap and Il1rapl,* although their direct contribution to NAD metabolism remains to be determined (Supplementary Fig. S3l).

Together, these findings indicate that partial loss of insulin signalling at the hippocampal vasculature enhances glutamatergic metabolism and mitochondrial ATP turnover, consistent with increased neuronal metabolism, while concurrently reducing hippocampal integration within anxiety-related brain networks, resulting in a robust anxiolytic phenotype.

### Hippocampal insulin resistance promotes long-term metabolic resilience

To examine the long-term trajectory of behavioural and neurometabolic adaptations in iNIRKO mice, we performed longitudinal assessments following initial phenotyping at 6 weeks post-tamoxifen induction. Female mice were subsequently exposed to either chronic corticosterone (CORT; 20mg/kg s.c., 20 days) or vehicle, and baseline behavioural and neuroimaging measures were repeated at 23 weeks post-tamoxifen (Fig. 4a). Chronic glucocorticoid excess impairs brain insulin signalling, predominantly through reduced IR/IRS phosphorylation and Akt activation^23,24^, providing an interventional approach to assess brain metabolic resilience and vulnerability to insulin dysregulation. Body weight did not differ between groups prior to CORT exposure, and CORT induced a comparable ∼10% increase in both genotypes, confirming the expected physiological response (Supplementary Fig. S4a-b).

**Fig. 4.**
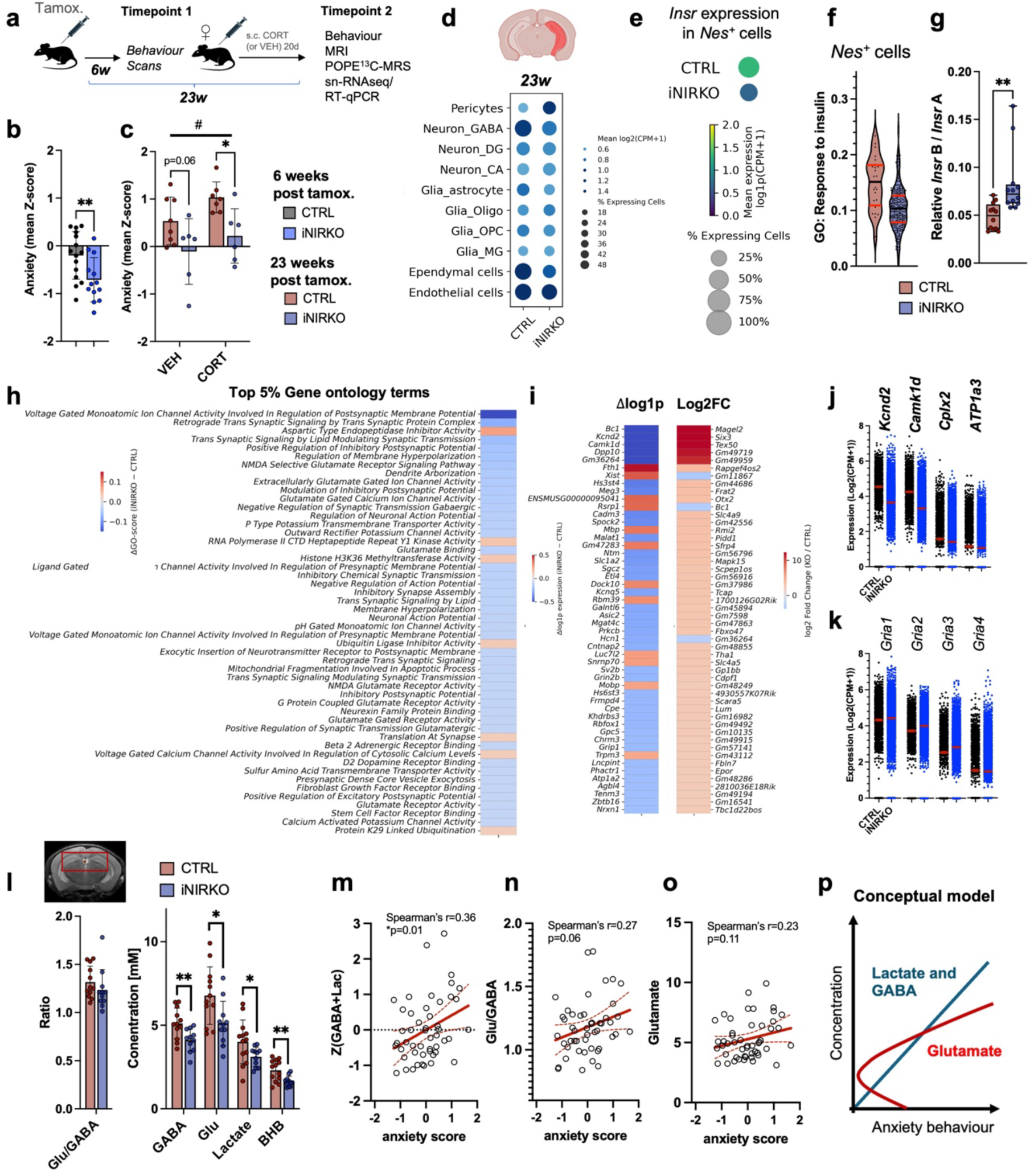
Hippocampal non-parenchymal insulin resistance stabilizes long-term metabolic and behavioural trajectories. a. Schematic of longitudinal behavioural and neuroimaging protocol. After the measurements presented above, i.e. at 6 weeks post tamoxifen induction (Fig.2 and 3), female mice underwent a second set of behavioural and neuroimaging measurements at 23 weeks post tamoxifen induction. CORT, corticosterone (20mg/kg); VEH, vehicle (saline). b. **,c.** Averaged anxiety (z-score) (**b**) 6 weeks and (**c**) 23 weeks post tamoxifen induction in iNIRKO (n_VEH_=6, n_CORT_=6) and CTRL(n_VEH_=8, n_CORT_=7) mice. (**b**) Student’s t-test, **p<0.005; (**c**) Two-way ANOVA, ^#^Genotype effect: F1,23 = 12.61, P = 0.0017; Treatment effect: F1,23 = 4.07, P = 0.055; followed by Bonferroni’s test, *p < 0.05. **d.** Dotplot of Insr gene expression across cell types in the whole hippocampus 23 weeks post tamoxifen induction in iNIRKO (n=10,296 cells, 6 pooled samples) and in CTRL (n=2,515 cells, 6 pooled samples) mice (all from VEH groups). **e.** Dotplot of Insr gene expression in Nestin positive (Nes^+^) cells in the whole hippocampus 23 weeks post tamoxifen induction. **f.** Violin plot of GOBP: Response to insulin (GO:0032868) gene scores for Nes^+^ cells. **g.** Relative Insr B and A isoform expression in dorsal hippocampus measured with qPCR. Student’s t-test, **p<0.005 **h.** Top 50 (5%) GO terms with highest difference in gene score (iNIRKO-CTRL) 23 weeks post tamoxifen induction **i.** Top 50 genes with highest change in gene expression (iNIRKO-CTRL) reported as difference (Δlog1p) or fold change (Log2FC) 23 weeks post tamoxifen induction. **j.,k.** Average expression of selected genes in excitatory neurons. Data are mean ± s.e.m **l.** POPE^13^C-MRS hippocampal voxel location and quantification of hippocampal metabolites before ^13^C-glucose infusion in CTRL (n=12) and iNIRKO (n=11) mice. Student’s t-test, *p<0.05, **p<0.005. **m.,n.,o.,** Correlation of hippocampal metabolite concentration with anxiety score (*p<0.05). Simple linear regression shown with 95% CI (red) **p.** Schematic of the relationship between observed metabolic changes and anxiety across CTRL and iNIRKO mice. All data are mean ± s.d. unless stated otherwise Tamox, tamoxifen; CORT, corticosterone; VEH, vehicle.

Chronic CORT treatment induced an anxiety-like phenotype in both groups, although iNIRKO mice consistently exhibited lower trait anxiety across conditions (Fig.4b-c). This behavioural pattern was accompanied by sustained reduction of *Insr*-related gene expression in Nestin-expressing hippocampal cells at 23 weeks (Fig. 4d-f), indicating stable long-term knock-out effect. Cell-type-resolved transcriptomics further confirmed that endothelial cells represent a major site of *Insr* enrichment in the mouse, and target of depletion in the iNIRKO model (Supplementary Fig. 4c-k). In addition, iNIRKO mice showed a higher Insr B-to-A isoform ratio (Fig. 4g), contrasting woith the relative enrichment of the A isoform reported in hyperinsulinaemia and Alzheimer’s disease, where reduces INSR-B abundance has been linked to impaired receptor signalling^25–27^. Transcriptomic profile remained highly stable over time, with persistent downregulation of excitability- and plasticity-related genes including *Kcnd2* and *Camk1d* (Fig.4h-k). Endothelial *Glut1* expression was consistently reduced in iNIRKO mice (Supplementary Fig.4l), pointing towards altered vascular glucose transport capacity and improved brain metabolism^28^.

Neurometabolic profiling using MRS revealed divergent longitudinal trajectories between genotypes. Whereas control mice exhibited age-associated increase in hippocampal lactate, GABA and glutamate, these changes were attenuated in iNIRKO mice (Fig.4l-p). Across animals and throughout the longitudinal study, lactate and GABA levels correlated with anxiety-related behaviour, mirroring their association with symptoms severity in humans, while glutamate concentrations alone showed weaker associations. These findings suggest that coordinated changes in neurotransmitter-related metabolites, rather than single metabolites, better reflect behavioural variation.

To further characterise glutamatergic metabolic flux, we examined hippocampal ^13^C-glucose tracing at 23 weeks. Control mice showed increased glutamate labelling with age, whereas iNIRKO mice remained stable across timepoints (Fig.5a-b), indicating an age-dependent divergence in glutamatergic glucose metabolism. We next examined lactate dynamics following ¹³C-glucose administration as an indirect readout of glycolysis-derived carbon flux. Unlabelled lactate production (ΔLac) was observed in both genotypes at 6 weeks which was reduced at 23 weeks, particularly in controls (Fig.5c), consistent with altered glycolysis in astrocytes over time. Notably, a similar effect was not observed for the ketone body BHB (ΔBHB; Supplementary Fig.S5a). To integrate these complementary metabolic dimensions, we derived a composite index combining glutamate labelling and lactate dynamics (¹³C-GluH4 × ΔLac/Lac), as a proxy for glycogen shunt activity because glycogen-derived carbon dilutes glutamate ^13^C labelling while contributing to the accumulation of unlabelled lactate. This exploratory metric was reduced in aged controls and correlated with anxiety-like behaviour (Fig.5d–e). Additional multimodal indices incorporating neurotransmitter concentrations and labelling patterns showed similar associations with behavioural variability (Supplementary Fig.S5b-d), supporting a convergent metabolic signature.

**Fig. 5.**
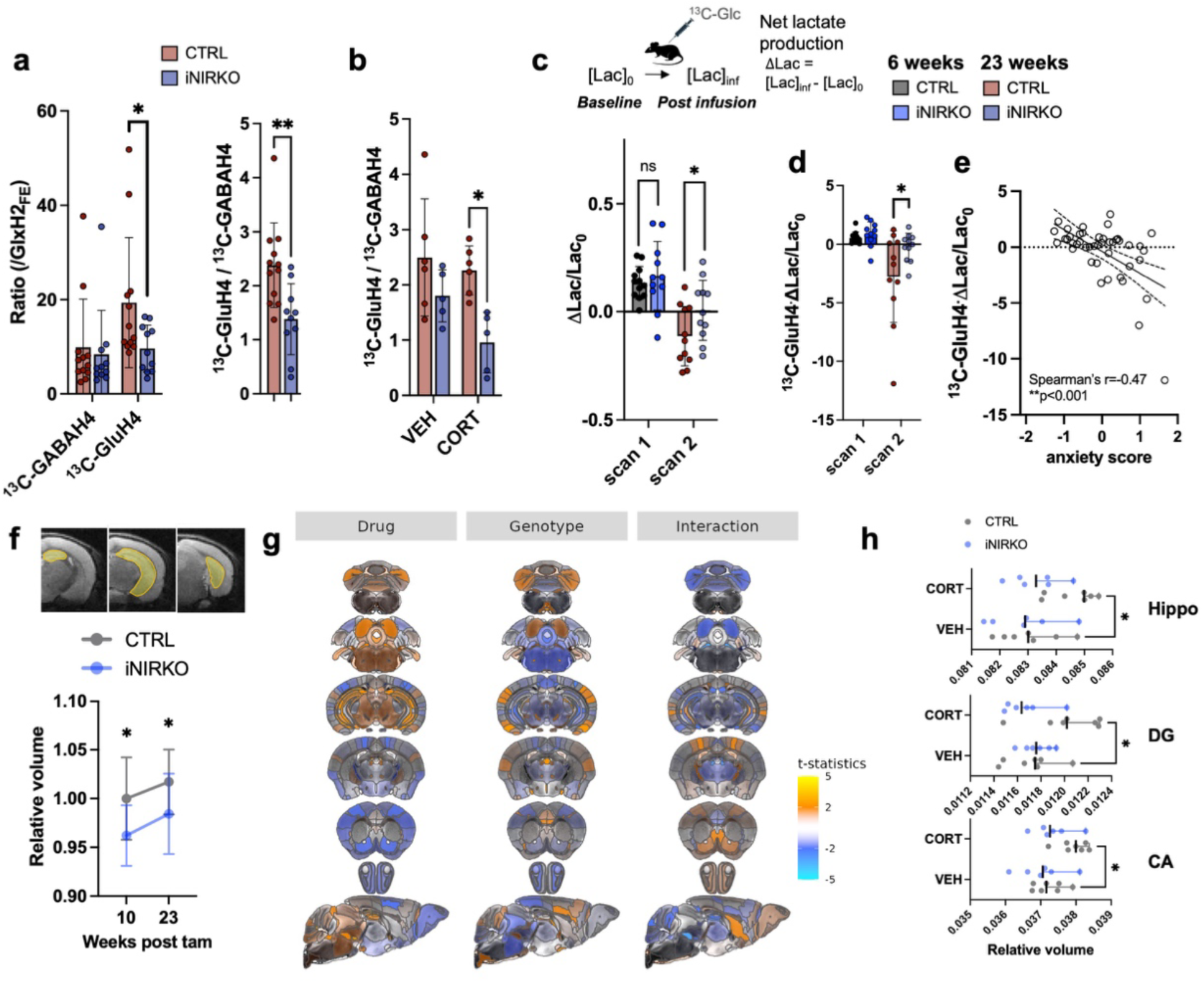
Hippocampal insulin resistance limits age-dependent glutamate-driven metabolic changes. **a.** Quantification of hippocampal relative metabolite labelling after ^13^C-glucose infusion 23 weeks post tamoxifen induction in CTRL (n=12) and iNIRKO (n=11) mice. No labelling in the lactate or BHB pools were detected. Labelling ratios of GABAH4 and GluH4 are reported relative to the fractional enrichment (FE) of GlxH2 (left) or relative to each-other (right). Student’s t-test, *p<0.05, **p<0.005. **b.** Relative labelling of GABAH4 and GluH4 following ^13^C-glucose infusion when considering treatment groups. Two-way ANOVA, Genotype effect: F1,18 = 11.29, P = 0.0035; Treatment effect: F1,18 = 3.27 P = 0.087 Bonferroni’s posthoc test *p<0.05 **c.** Relative difference in total (unlabelled) lactate concentration before and after ^13^C-glucose infusion at 6 and 23 weeks post tamoxifen induction. Student’s t-test, *p<0.05. **d.** Group differences in composite metabolic marker ^13^C-GluH4*Δlactate/lactate at 6 and 23 weeks post tamoxifen induction. Student’s t-test, *p<0.05. **e.** Relationship of ^13^C-GluH4*Δlactate/lactate with anxiety. Data from both 6 and 23 weeks timepoints. Correlation (**p <0.001) and simple linear regression fit (*β*=-0.98) shown with 95% CI. **f.** Longitudinal volume change of the hippocampus in CTRL (N=13) and iNIRKO (N=15) mice measured from manual segmentation of anatomical MRI. Two-way ANOVA, Genotype effect: F1,25 = 8.70, P = 0.0066; Time effect: F1,25 = 6.55 P = 0.017 Bonferroni’s posthoc test *p<0.05. **g.** Voxel-wise maps for selected slices of the mouse brain showing statistically significant volume differences (FDR q<0.05) as an effect of drug (CORT) or genotype, as well as interaction (genotype x drug). **h.** Volume differences of hippocampal subfields. Including hippocampal formation (Treatment effect: F1,22 = 6.02, P = 0.02), dentate gyrus (DG; Interaction: F1,22 = 5.72, P = 0.03) and cornu ammonis (CA; Treatment effect: F1,22 = 6.23, P = 0.02). Two-way ANOVA, Bonferroni’s posthoc test *p<0.05.

Transcriptomic analysis revealed enrichment of genes involved in astrocytic glutamate handling and inhibitory signalling pathways (Supplementary Fig.S5e-j), although classical excitotoxicity signatures were not prominent. Instead, increased expression of glutamate transport and calcium signalling-related genes (including *Slc1a2*) was observed in controls, suggesting altered regulation of glutamatergic homeostasis rather than overt excitotoxic damage.

Finally, iNIRKO mice exhibited reduced hippocampal volume at both timepoints, whereas CORT induced volume increases that were attenuated in iNIRKO mice (Fig.5f-h), particularly in the dentate gyrus (treatment x genotype interaction). This structural effect coincided with reduced vascular *Glut1* expression and inflammatory signalling-related genes (Supplementary Fig.5k-l), suggesting altered neurovascular responsiveness to chronic stress.

Together, these findings indicate that hippocampal endothelial insulin receptor reduction modifies the trajectory of age- and stress-related metabolic and structural changes, and is associated with a more stable neurometabolic profile over time.

## Discussion

The mechanisms by which insulin signalling in the brain influences mood remain incompletely understood, and brain insulin resistance is often considered to be a uniformly maladaptive state. Here, we show that insulin signalling exerts cell-type-specific effects within the hippocampus. In particular, reduced insulin action at the neurovascular interface enhances neuronal metabolism and attenuates anxiety-like behaviour in mice. In parallel, multimodal neuroimaging in humans identifies a hippocampal signature - elevated lactate and GABA with increased hippocampus-DMN connectivity - that tracks glycaemic variation and mood symptoms. Together, these findings suggest that brain insulin resistance is not a unitary deficit but a context-dependent process linking metabolism to affective state.

Our findings challenge the prevailing view that brain insulin resistance is uniformly detrimental. We show that insulin resistance is cell-type-specific, with distinct consequences depending on whether it arises in parenchymal or non-parenchymal compartments. Importantly, this manipulation differs fundamentally from previous models. Rather than producing severe neuronal or whole-brain insulin receptor deficiency, our model captures a mild (20%), hippocampus-restricted reduction in *Insr* expression that preferentially affects the neurovascular interface. As a result, neuronal insulin responsiveness is preserved, and may even be enhanced, indicating that the physiological consequences of insulin resistance depend on both its cellular localisation and the extend of signalling impairment. Genetic ablation of the insulin receptor (*Insr*) in the brain has consistently been associated with impaired synaptic plasticity, altered neurotransmission and depressive-like phenotypes across a range of region- and cell-type-specific approaches, including whole-brain^21,29–31^, hypothalamic^32–37^, hippocampal^38,39^ and amygdalar^38^ deletions, as well neuronal^32–34,39^, glial^40–44^ and whole body vascular^45^ targeting. In contrast, we show that selective attenuation of insulin signalling in non-parenchymal cells produced the opposite profile—enhanced metabolism and reduced anxiety-like behaviour. This apparent paradox likely reflects a fundamental distinction between reducing insulin responsiveness within neurons and reducing insulin responsiveness at the blood-brain barrier. The blood–brain barrier actively regulates brain insulin action^26,46,47^, functioning as a key gatekeeper of insulin transport and signalling^48^ – a process disrupted under metabolic stress such as diet-induced pathology^49,50^. Our findings suggest that the consequences of brain insulin resistance depend critically on where it occurs. Whereas neuronal insulin resistance impairs insulin-dependent cellular processes, reduced insulin responsiveness at the neurovascular interface may serve an adaptive gatekeeper function, limiting excessive metabolic exposure to the parenchyma. The observation that hippocampal lactate and GABA track affective symptoms in humans and anxiety-related behaviour in mice supports the translational relevance of this interpretation.

We propose a working model (Fig.6) in which reduced insulin signalling at the neurovascular interface may limit excessive insulin and glucose entry into the brain^51,52^, thereby preserving metabolic homeostasis under conditions of metabolic load (allostasis^53^). Non-parenchymal insulin resistance may consequently reshape hippocampal metabolism and circuit function through compensatory increases in neuronal insulin sensitivity. Consistent with this interpretation, iNIRKO mice exhibited elevated oxidative flux and ATP turnover, altered neurotransmitter profiles, reduced hippocampal functional connectivity and decreased expression of genes regulating neuronal excitability, including *Kcnd2*^54–57^. Together, these changes are consistent with altered excitatory-inhibitory balance and reduced large-scale synchrony. A metabolically engaged but less synchronised hippocampus may favour exploratory over threat-driven response^58^, in line with the anxiolytic phenotype observed in our model. This state may be associated with reduced reliance on astrocytic glutamate recycling^59^ and GABAergic inhibition^60^, potentially lowering allostatic burden on these cell types^53^. Although these mechanisms remain to be directly tested, they are consistent with the observed transcriptional, metabolic and behavioural adaptations. Together, our findings support a model in which the neurovascular interface acts as a regulator of hippocampal metabolic state, linking systemic metabolic signals to circuit dynamics and behaviour.

**Fig. 6.**
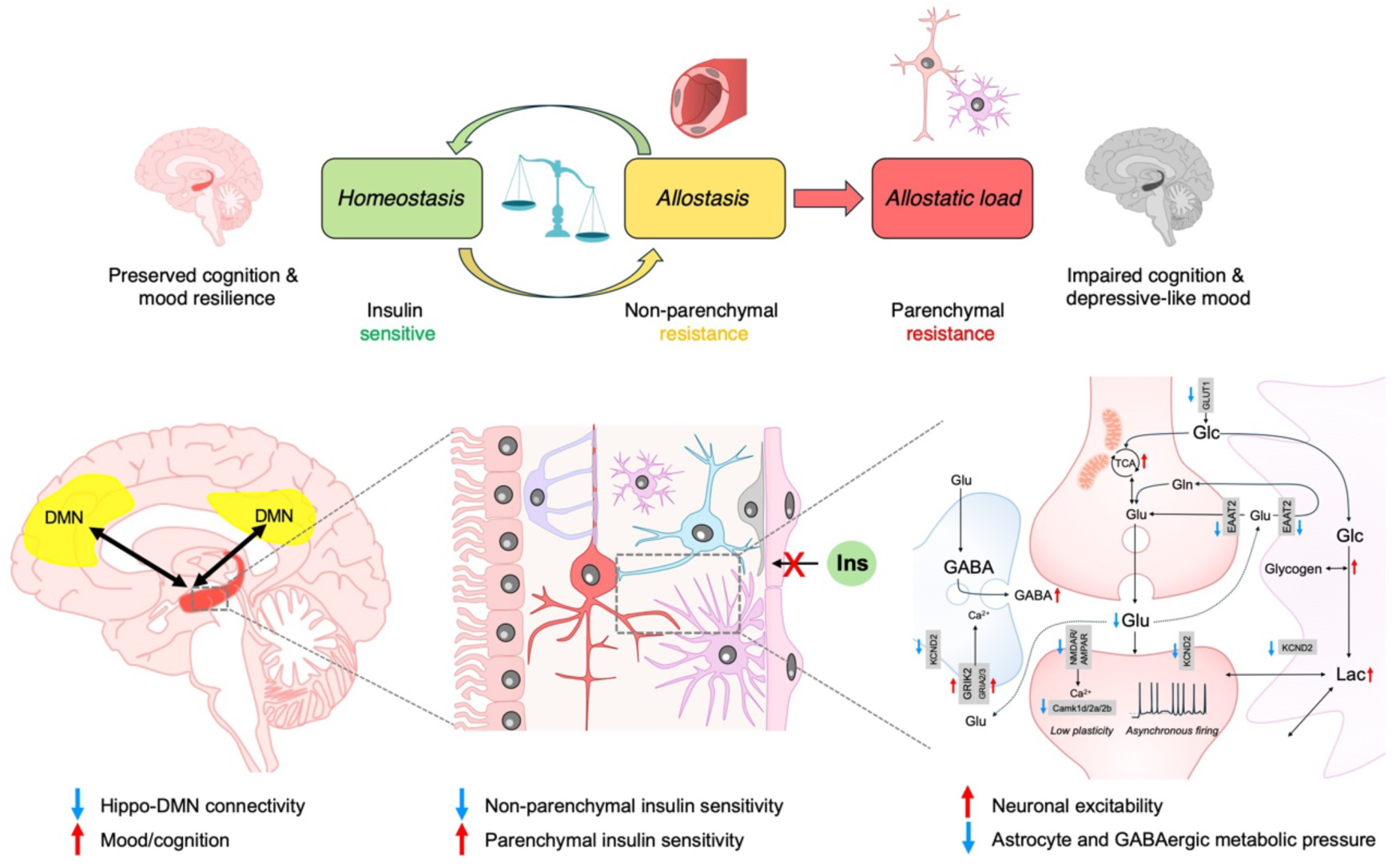
Summary scheme. **(top**) Successive forms of brain insulin resistance differentially regulate hippocampal function and mood: neuronal resistance is detrimental, whereas vascular resistance acts as a protective (allostatic) gatekeeper that shields the parenchyma from excessive metabolic pressure (allostatic load). (**bottom**) Schematic of metabolic changes in iNIRKO mice. Deletion of insulin receptor in non-parenchymal cells upregulates insulin sensitivity in neuron and glial cells, increasing neuronal excitability and metabolic health, while reducing hippocampal network connectivity within the default-mode network (DMN), ultimately improving mood.

The hippocampus emerges as a key node in this process. Across species, we identify a hippocampal metabolic and functional signature that is tightly coupled to glycaemic state yet dissociable from global adiposity. The regional specificity of this signature, together with the effects of hippocampal-targeted manipulation in mice, suggests that insulin-related modulation of brain function is preferentially expressed within hippocampal circuits. Given the hippocampus’ role in integrating metabolic signals^61–64^ and regulating large-scale network dynamics, including connectivity with HPA-axis and the default mode network^65^, these findings position it as a critical interface between systemic metabolism and affective regulation. Insulin modulates hippocampal–DMN connectivity^66,67^, a network shown to be affected in depression^68–70^. The hippocampus is thought to have a functional gradient^71^, with the posterior (dorsal in rodents) part classically associated with cognition and spatial memory, while the anterior (ventral in rodents) part regulates mood, emotion, and stress responses^72–74^. Consistently, we show that, in humans, anterior hippocampal–DMN connectivity most strongly correlated with anxiety, whereas in our mouse model, dorsal hippocampus effects were more pronounced, reflecting region-specific *Insr* knock-out and enhanced sensitivity within the dorsal neurogenic niche.

Our results also define a candidate neuroimaging signature of altered BIR, currently lacking in the field^18,75^. By combining measures of functional connectivity with metabolic readouts, including GABA and lactate, we identify a translational phenotype that is conserved across mice and humans and preferentially associated with affective symptoms. While DMN connectivity has been recently proposed as a candidate biomarker for measuring BIR^18^, metabolic measures such as MRS provide molecular insight allowing additional mechanistic interpretation and showed more specific association with affective state. Although further validation is required, our study provides a framework for *in vivo* assessment of brain insulin sensitivity and for stratifying individuals according to neurometabolic state.

Several limitations should be considered. The human findings were obtained in a relatively small cohort of unmedicated individuals with depression and therefore require replication in larger independent samples. Although the human data are necessarily observational, the parallel findings in the iNIRKO model provide experimental support for a mechanistic link between insulin signalling, brain metabolism and affective behaviour. The mouse model captures a partial and regionally restricted reduction in insulin receptor expression, and although it identifies neurovascular insulin signalling as a key regulator of hippocampal metabolism and behaviour, the downstream cellular mechanisms linking these effects remain to be fully elucidated.

## Materials and Methods

### Human participants

#### Study design

All experiments were conducted in accordance with the principles of the Declaration of Helsinki and were approved by the Health Research Authority (HRA) and Health and Care Research Wales (HCRW) Research Ethics Committee (REF: 22/WA/0243). The cross-sectional study was preregistered on the Open Science Framework (OSF; https://doi.org/10.17605/OSF.IO/4YN2V). Written informed consent was obtained from all participants, comprising 9 patients with depression (3 females, 6, males, age 38±12 years, mean±s.d.) and 12 healthy volunteers (6 females, 6, males, age 34±14 years, mean±s.d.). All patients had a prior clinical diagnosis of depression and reported ongoing depressive symptoms at the time of study participation. Diagnosis was confirmed during the screening interview using the Structured Clinical Interview for DSM-5 (SCID-5)^76^. Healthy controls were recruited from the general population. Exclusion criteria for patients included a current or past diagnosis of bipolar disorder, schizophrenia or emotionally unstable personality disorder, as well as a clinically significant risk of suicidal behaviour. For healthy controls, exclusion criteria included any past or present DSM-5-defined mental health condition. For both groups, additional exclusion criteria were contraindications to magnetic resonance imaging (MRI), a history of severe claustrophobia, pregnancy, or breastfeeding, the use of any medication or substance that could influence brain function, such as benzodiazepines, recreational drugs, or excessive alcohol consumption, as well as medications for metabolic conditions, including metformin, statins, and antihypertensive drugs. Although this was not a formal exclusion criterion, all patients were not taking antidepressant medication at the time of the study visit. Current and past mental health history was assessed for all participants using the SCID-5 during screening.

All participants underwent a single testing session at the Functional Magnetic Resonance Imaging of the Brain (FMRIB) building in Oxford which included an MRI scan, body measurements and cognitive testing detailed below.

#### Brain imaging

Brain MRI and MRS acquisitions were conducted on a Magnetom 7 T Plus scanner (Siemens Healthineers, Erlangen, Germany) with a Nova Medical 8-transmit/32-receive head coil. T1-weighted images were acquired from each subject using the following parameters: TE/TR = 3.31ms/2.6s; flip angle = 5°; Field of view (FOV) = 224 mm; voxel size= 0.7x0.7x0.7 mm^3^. After this anatomical acquisition, resting-state functional MRI (rs-fMRI) data was acquired for 10 min using an EPI multiband sequence^77,78^ with following parameter: TE= 12.8ms; TR=1087ms; FOV=216mm; slices= 64; voxel size = 2x2x2 mm^3^; multiband acceleration factor = 2. During the resting-state session, participants were asked to keep their eyes closed but try not to fall asleep. Anatomical images were used to place the ^1^H-MRS voxel in the left hippocampus (voxel size = 16x16x35 mm^3^) and dorsal anterior cingulate cortex (dACC; voxel size = 16x20x35 mm^3^). ^1^H-MRS was acquired with MEGA-sLASER^79^ (CMRR Spectroscopy Package, University of Minnesota) for the measurement of GABA (GABA+) at 3.0 ppm and for lactate at 1.31ppm, using following parameters: data points = 2048; Averages = 64; TR = 4s; TE_GABA_=78ms, TE_LAC_=100ms; editing pulse bandwidth = 130 Hz; editing pulse offset (ON/OFF) Offset_GABA_ = 1.90/7.50ppm and Offset_Lactate_ = 4.10/5.20ppm. Shimming was performed with Fastmap^80^ and acquisitions included VAPOR^81^ and Outer Volume Suppression (OVS) (70mm slice with 3mm offset from voxel). Unsuppressed water acquisitions were performed with the same region-specific parameters for the quantification.

#### Brain Imaging analysis

Segmentation of the hippocampus was performed using FSL-FIRST^82^ of the FSL toolbox^83^ and volumes were measured using fslstats. Edited MRS spectra were processed in jMRUIv6.0 (www.jmrui.eu^84^), including phase and frequency correction, followed by (ON-OFF) subtraction. Metabolite concentrations in the hippocampus and dACC were quantified using the QUEST algorithm^85^, with a basis set derived from phantom-acquired metabolite signals, using the unsuppressed water signal as internal reference assuming 80% tissue water content. Resting-state fMRI data was pre-processed using FEAT^86^ for motion correction (MCFLIRT)^87^, high-pass filtering and fieldmap correction. Data was then further denoised from 20 MELODIC^88^ ICA components with FMRIB’s ICA-based Xnoiseifier (FIX)^9^ using a pretrained classifier (HCP7T_hp200) and then registered to standard space. Anterior and Posterior Hippocampus were selected *a priori* as seeds for whole-brain voxelwise analyses, and defined from the HarvardOxford Atlas following previously reported anatomical subdivisions^91^. Group-level nonparametric permutation testing was performed with *randomise*^92^,using age and sex as covariates.

#### Body measurements

After the scan participant underwent a blood test for measuring systemic insulin resistance (HbA1c), followed by body measurements using a BF511 Body Composition Monitor (OMRON Healthcare), which included BMI, % body fat, % body muscle, resting metabolic rate and visceral fat. HbA1c values were analysed by the Biochemistry laboratory at the Oxford University Hospitals (OUH).

#### Cognition and mood assessment

Depression, anxiety and related symptom dimensions were assessed using a set of self-report questionnaires—including the Quick Inventory of Depressive Symptomatology – Self-Report (QIDS-SR)^93^ for depressive symptom severity, the Hospital Anxiety and Depression Scale-Anxiety subscale (HADS-A)^94^ for anxiety symptoms, the Snaith-Hamilton Pleasure Scale (SHAPS)^78^ for anhedonia, and the Apathy Motivation Index (AMI) ^96^ for apathy and motivational deficits.

Cognition was assessed with the Montreal Cognitive Assessment (MoCA) administered in person^97^, and digitally using the Oxford Cognitive Testing Portal (OCTAL) battery^98^, a remote, browser-based cognitive assessment platform designed for unsupervised administration on personal devices (https://octalportal.com). OCTAL comprises a suite of tasks measuring memory, attention, executive function and visuospatial processing, developed using PsychoJS and hosted on Pavlovia.org. The battery has been validated against the Addenbrooke’s Cognitive Examination-III (ACE-III) in a memory-clinic cohort including patients with Alzheimer’s disease, mild cognitive impairment and subjective cognitive decline as well as age-matched healthy controls, where a 20-minute OCTAL composite achieved an area under the curve of 0.97 for distinguishing neurodegenerative from subjective impairment, even better than the ACE-III. In a healthy population with more than 1100 participants across lifespan, OCTAL captured domain-specific age-related cognitive trajectories with high sensitivity, and all principal metrics demonstrated good-to-excellent 6-month test–retest reliability^98^. The version administered in the present study included the Trail Making Test (TMT)^99^, assessing processing speed and cognitive flexibility; the Digit Symbol Substitution Test (DSST)^100^, measuring attention and processing speed; the freestyle Corsi block-tapping test (CORSI)^101^ testing visuospatial short-term memory; the Object Memory Task (OMT)^102^, assessing short-term memory for object identity and location; the Rey-Osterrieth Complex Figure (ROCF)^103^ test, a digital drag-and-drop adaptation measuring visuo-constructive ability and visuospatial episodic memory; the Objects In Scene (OIS) test^104^, assessing associative memory for objects within naturalistic scenes at short-term and long-term delays; and a verbal memory wordlist recall task (ALF) adapted from the Rey Auditory Verbal Learning Test and accelerated long-term forgetting paradigms, comprising immediate free recall and recognition followed by 30-minute delayed recall and recognition. All tasks were completed on a tablet via the OCTAL platform; task-specific attention checks were embedded to ensure data quality. Full task descriptions, scoring procedures and validation details are provided in Zhao et al.^98^

Principal component analysis (PCA) was performed on 34 behavioural variables derived from mood questionnaires and cognitive task performance across all participants. Prior to analysis, variables were z-score standardized to ensure comparable scaling across measures. Cognitive variables were oriented such that higher z-scores reflected worse performance (e.g., slower reaction times or larger error magnitudes), ensuring consistent interpretation across measures. To reduce dimensionality and derive composite measures of cognitive performance, domain-level scores were calculated as the mean of standardized z-scores from individual task measures. The following cognitive domains were defined: Executive function (TMT_A, DSST_RT), Short-term memory (OMT_ProportionCorrect_3item_4secs, AbsoluteError_3item_4secs, OMT_Misbinding, OMT_Guessing, OMT_imprecision, CORSI_3itemError, WM_nCorrect_freerecall_1, OIS_ImmediateObjectAccuracy, OIS_ImmediatLocationError), Long-term verbal memory (WM_propcorrect_recognise_delayed, WM_nCorrect_freerecall_delayed) and Long-term visuospatial memory (OIS_DelayedObjectAccuracy, OIS_DelayedLocationError). The number of principal components retained was determined using parallel analysis with 1,000 Monte Carlo simulations. Component scores were calculated for each participant and used in subsequent analyses. Group differences in component scores were assessed using unpaired Student’s t test. Associations between behavioural variables, cognitive domain scores, neuroimaging metrics and HbA1c were assessed using Spearman correlation analyses and p-values were not corrected for multiple comparison but reported for all comparisons.

### Animals

Animals were all housed in standard IVC cages in a normal 12h day-light cycle environment at a temperature of 20-24°C and humidity of 46-65%. Animals had *ad libitum* access to standard rodent chow diet and water. Animal care and experimental procedures were approved by the University of Oxford local ethical review committee, and all experiments were carried out with approval by the UK Home Office, under the Animals (Scientific Procedures) Act 1986, and in compliance with the ARRIVE (Animal Research: Reporting *in vivo* Experiments) guidelines.

### iNIRKO breeding and generation

The iNIRKO mice (*Insr^fl^*^/fl^; *Nes*-Cre/ERT2^+/-^) were generated from *Insr^fl^*^/fl^ (B6.129S4(FVB)-*Insr^tm1Khn^*/J; strain 006955) and Nes-Cre/ERT2^+/-^ (C57BL/6-Tg(Nes-Cre/ERT2)KEisc/J; strain 016261) obtained from the Jackson Laboratory (Farmington, US). The breeding was set up in-house to produce the iNIRKO mice and their control littermates (CTRL; *Insr^fl^*^/fl^; *Nes-*Cre/ERT2^-/-^). The induction of insulin receptor recombination was achieved using a 5-days oral administration of a 100 ul dose of tamoxifen (Sigma-Aldrich, CAS# 10540-29-1) solution in corn oil (20 mg/ml) to reach a dose of (75 mg/kg body weight).

The inducible Nestin-CreERT2 line is widely used to target neural stem and progenitor cells in the adult mouse brain. The strain used here^105^ exhibits low recombination efficiency in the dentate gyrus, contributing minimally to mature neuronal populations (<1% in the SGZ/SVZ) and showing limited ectopic labelling^106^. However, the cellular specificity of the recombination remains difficult to predict, as insulin signalling in neural stem cells can influence downstream neurogenesis in an age-dependent manner^107^. Importantly, Nestin expression is not restricted to neural progenitors and has also been reported in endothelial progenitor (EPC) and mature endothelial cells (EC)^108^ (Supplementary Fig.S2). Although the second intronic enhancer of the Nestin gene is thought to confer central nervous system specificity and limit recombination in EPCs^109^, the extend of recombination within brain vascular populations remains uncertain. Furthermore, recombination patterns in Nestin-CreERT2 lines vary across strains^106^ and tamoxifen administration protocols (e.g. dose, timing and control group^107^), which potentially impacts neurogenesis^110^. In addition, labelling of non-neurogenic cell types, including ependymal cells, has been reported^111^. To resolve this, we performed single-nucleus RNA sequencing to assess the effective cellular distribution of *Insr* deletion in the iNIRKO model under our current tamoxifen protocol. This analysis revealed that the depletion of *Insr* expression was restricted to the dorsal hippocampus, in line with a neurogenesis-driven effect, but effectively only visible among vascular cells (see Results).

### Behaviour testing

Anxiety tests assessed the approach-avoidance conflicts of mice when exposed to an anxiogenic environment. These tests capitalize on animal’s innate fear of bright-lit, open or elevated spaces^112^. The details of the individual tests and their conversion into an anxiety score are described below. Additionally, the number of boli present on the floor at the end of the experiment were counted in each test and averaged for each mouse to get a defecation score. Defecation in rodents following a stressful stimulus is a result of complex physiological and hormonal changes triggered by the body’s stress response^113^. Finally, a control experiment assessed the locomotor activity in a non-anxiogenic environment.

#### Elevated Plus Maze (EPM) test

The EPM test was performed using a 50 cm-high platform placed in a dimly lit room. The open arms (35 x 6 cm^2^) were perpendicular to the closed arms (35 x 6 cm^2^) with a 6 x 6 cm^2^ central region in between. Illuminance values were measured at 5-10 lux in the closed arms and 60-75 lux in the open arms. Mice were placed in the central zone and automatically tracked for 600 s (ANY-maze software, Stoelting Co.). The time spent in the closed and open arms, the number of entries in the open arms and the immobility were calculated from the tracking.

#### Light-Dark Box (LDB) test

The LDB test was performed with an arena placed inside a sound-attenuating cubicle and was composed of a bright (280 lux) open compartment (46.5 x 21 x 21 cm^3^) and a small black (2 lux) covered compartment (21 x 16 x 16 cm^3^) with a small connecting opening connecting the two compartment (3 x 2.7 cm^2^). Animals were placed in the dark compartment and tracked for 600 s (ANY-maze software, Stoelting Co.). The distance travelled in the lit compartment as well as the number of entries and the latency to first enter that compartment were calculated from the tracking.

#### Aversive Open Field (AOF) test

the AOF test was performed in an enclosed white cylindrical chamber (30 cm radius) with bright light (475-500 lux). Mice were initially placed in the side of the chamber, and the movement was automatically tracked for 600 s (ANY-maze software, Stoelting Co.). A centre zone (10 cm radius) was delineated a posteriori on the videos for the estimation of the activity. Key parameters were distance travelled in the centre zone and the number of entries done in the centre zone.

#### Locomotor activity (LMA) monitoring

Monitoring of locomotion was performed in an unfamiliar but low-anxiety environment, i.e. a transparent Plexiglas box (42 x 22 x 20 cm) lined with sawdust and covered with a perforated Plexiglas lid. Animals were placed in each cage individually in a dimly lit room [∼60 lux] and were allowed to explore freely for 3600 s while being monitored. Monitoring was done using Photo Beam Activity Hardware and Software (Photobeam Frame software, San Diego Instruments). Number of beam breaks was automatically recorded in 5 sec bins across the box for horizontal and vertical (rearing) motion of the mouse.

#### Anxiety score

An anxiety score was computed for each mouse based on the different metric acquired during behavioural testing. To do so, an average z-score was computed for each of the anxiety tests, which were then averaged to get a single z-score of anxiety including outputs from the EPM, LDB and AOF tests. More specifically, four z-scores were calculated: Z_EPM_=(Z(Time in closed arm)-Z(Time in open arms)-Z(Number of entries in open arms))/3, Z_LDB_=Z(Latency to first enter lit compartment)-Z(Distance in lit compartment)-Z(Number of entries in lit compartment))/3, Z_AOF_=(-Z(Distance travelled in centre zone)-Z(Number of entries in centre zone))/2, Z_immo_=(Z(time immobile in EPM)+Z(time immobile in AOF))/2. All four z-scores were then averaged to compute the final anxiety score = (Z_EPM_ + Z_LDB_ + Z_AOF_ + Z_immo_)/4.

### *In vivo* Magnetic Resonance Imaging (MRI)

MRI scans included magnetic resonance spectroscopy (MRS), resting-state functional MRI (rs-fMRI) and structural imaging, and were acquired on a 7T (70/20) BioSpec MRI scanner (Bruker, Ettlingen, DE) equipped with either a transmit (Tx) 86 mm volume coil and a receiver (Rx) ^1^H cryoprobe or a 10 mm dual tuned ^31^P/^1^H (Tx/Rx) coil (PulseTeq Ltd, Chobham, UK) using Paravision 360.1.1. Mouse anaesthesia was induced with a bolus of 4% isoflurane mixed with air, followed by a bolus of medetomidine in saline (0.2 mg/kg, 0.1 mg/ml, s.c.) and a continuous infusion (0.4 mg/kg/h, s.c.) with minimal isoflurane administration (0.2-0.3% in air containing 35% O_2_). Monitoring of the breathing rate and rectal temperature was achieved during the entire scans using a small animal monitoring system (SA Instruments Inc., New York, USA). No breathing or temperature group differences were observed across all in vivo MRI experiments as shown in Supplementary table S1. At the end of the experiment, anaesthesia was reversed using single bolus of atipamezole (1 mg/kg) solution (0.5 mg/ml). Details of the different acquisitions and analysis for each MRI modality are presented below:

### In vivo proton-observed proton edited ^13^C- Magnetic Resonance Spectroscopy (POPE^13^C-MRS)

*In vivo* ^13^C-glucose isotopic labelling experiments were performed with POPE^13^C-MRS, as previously described^22^. Briefly, baseline GABA and lactate POPE^13^C-MRS scan (t0) were acquired from the hippocampus (voxel size: 6.5 x 2.5 x 3 mm^3^) for 32min, followed by a constant infusion of 99% uniformly ^13^C-labelled glucose ([U-^13^C_6_]-Glc, 20% w/v) solution in the animal’s flank (s.c., 2.83 mL/kg bolus, followed by 10 mL/kg/h). After the first 30min of infusion, a second MRS scan (tf) was acquired for 51 min during the ongoing infusion. The MRS voxel placement was aided by a set of anatomical T_2_-weighted localizer (FLASH, TE/TR= 3ms/15ms, 2 averages x 1 repetition). Shimming (MAPSHIM) allowed for a water FWHM below 15Hz in each subject. POPE^13^C-MRS (Npoints=2048; AcquisitionBW=9.82ppm; TR=3s; TE_GABA_=69.5ms, TE_LAC_=106ms; Offset_GABA_= 1.9ppm, Offset_Lac_=4.1ppm; t0 (repetitions x averages) = 5 x 32, tf (repetitions x averages) = 8 x 32) was acquired with OVS and water suppression (VAPOR). Spectra transients were post-processed using the jMRUIv6.0 routine (phase and frequency correction, subtraction and NAA signal removal) and then analysed in QUEST^85^ using a homemade basis set including each metabolite component and associated ^13^C-satellites. Metabolite concentrations were then estimated using the unsuppressed water signal as internal reference, after correcting for differences in tCr peak between t_0_ and t_f_, and metabolite labelling concentrations were referenced to GlxH2_FE_ to correct for differences in glucose infusion efficiency. Additionally, the t_0_ scans were used to quantify baseline metabolite concentrations of GABA, glutamate, Glx, Lactate and BHB.

#### Structural MRI

A set of T2-weighted 150um isotropic MRI images were acquired (TE_eff_=22ms, Rarefactor=3, TR=6.1s, field of view = 12 × 8.1 x 9.6 mm, and matrix size = 256 x 256 x 64). Additionally, whole brain volumetric analysis was done using an automated registration into a consensus average space as previously described^114^. Briefly, registration onto the consensus space was done using Pydpiper, then MAGeT Brain algorithm^115^ was used to register a pre-segmented atlas^116^ containing 183 bilateral brain regions onto each image, resulting in a single volume measurement for each brain structure. These structures were then fitted with a mixed-effects model and multiple comparisons were corrected using false discovery rate (FDR)^117^.

#### Resting-state fMRI (rs-fMRI)

Resting-state functional MRI (rs-fMRI) was acquired during the POPE^13^C-MRS experiments immediately before the second (tf) MRS acquisition, and 20 min after the start of the glucose infusion. Anatomical T_2_-weighted imaging was acquired (RARE, TE_eff_/TR= 33ms/2500ms, averages=6, RARE-factor=8, matrix size=256x256x64, FOV=15x10.8mm, spatial resolution=100x100um, slice thickness=500um) prior to the fMRI acquisition for registration purpose during the post-processing steps. Blood oxygenation-dependent (BOLD) resting-state fMRI time series were acquired with an Gradient-Echo Echo Planar imaging (GE-EPI) sequence (TE/TR: 0.345ms/1s; Effective EPI bandwidth = 2898.55 Hz; in plane resolution = 0.2 x 0.2 mm^2^, number of slices=20; slice thickness =0.5 mm, slice gap=0mm; 800 volumes) leading to a 13.3min acquisition.

The fMRI data was pre-processed using the RABIES pipeline^118^, performing registrations to a data-driven study template, and implementing AROMA^119^ for ICA-based denoising. ROIs were selected *a priori* based on the DSURQE atlas^120^. The hippocampus see was divided into dorsal and ventral ROIs in alignment with previous literature on cross-species correspondence of hippocampal anatomy^71^; separation into dorsal and ventral hippocampal ROIs was performed manually to match MRS voxel placement. Functional connectivity values are given as Pearson correlation coefficients, which were Fisher r-to-z transformed before statistical comparisons with Student’s t-tests.

#### Steady-state ^31^P-MRS

^31^P-MRS was conducted as previously described^121^, using a dual tuned ^31^P/^1^H surface Tx/Rx coil (PulseTeq Ltd, Chobham, UK). MRS voxel placement was achieved following acquisition of a set of anatomical T_2_-weighted images acquired in axial, sagittal and coronal orientations (TurboRARE, TE/TR= 11ms/2632.7ms, RARE-factor=8) in a 140uL voxel (8x5x3.5mm^3^) englobing bilateral whole hippocampi. Shimming (MAPSHIM) was performed in the voxel to reach a water linewidth_50%_ of 26±2Hz, followed by the acquisition using 3D-ISIS (Npoints=1024, AcquisitionBW=40ppm, TR=4s, Averages/Repetitions=64x18), leading to an overall scanning time of ∼2h per mouse. Spectra were processed (phasing, B0-drift correction, averaging) in jMRUI and analysed with AMARES^84^, using Lorentzian line-shape and constrained frequency, linewidth and amplitude for each component (PCr, γATP, αATP, βATP, P_i_, PE, PCho, GPC, GPE, NADtot, NAD^+^). Relative Cramer-Rao Lower Bounds (CRLB) were calculated by dividing the fit error (SD) obtained from AMARES quantification over the signal amplitude^122^. The SNR_PCr_ was measured using MATLAB and was comparable across groups (8.7±2.2 for CTRLs and 8.5±1.3 for iNIRKO). The pH was determined from the chemical shift difference between Pi and PCr for each mouse. No difference across groups was observed in the animal’s physiology during the scans (supplementary table S1)

#### Saturation-transfer ^31^P-MRS

Saturation transfer ^31^P-MRS experiments were performed as previously described^121^, using a 120uL spectroscopic voxel (8x5x3mm^3^) placed in the whole bilateral hippocampi. To limit the effect of contaminating signal from non-hippocampal adjacent tissues, Outer Volume Suppression module was included (5mm thick slices with a 0.5mm gap from each voxel side). The water peak Full Width at Half Maximum (FWHM) was 24±2Hz after shimming with MAPSHIM. Progressive saturation transfer experiment was done using a BISTRO 3D-ISIS sequence (Npoints=1024, AcquisitionBW=40 ppm, TR=6s)^123^, consisting of eight 40ms HS2 pulses (BW=100Hz) of variable RF amplitude (scaling factors: 0.02, 0.04, 0.07, 0.14, 0.27, 0.49, 0.82, 1) at -2.50 ppm on the yATP resonance. Saturation time was acquired in random order at 5 different values (336, 672, 1’009, 2’018 and 4’710 ms) and control spectrum was acquired with a mirrored saturation at +2.50ppm. Each saturation transfer scan lasted ∼2h per subject. After spectra post-processing in jMRUI, the PCr signals were fitted using a single Lorentzian line-shape in AMARES. A mono-exponential function was fitted (MATLAB) to the relative PCr signal (M_PCr_) decay as a function of saturation time (t_sat_) using following equation: M_PCr_(t_sat_)/M_PCr(_0) = (1-k*T1)+k*T1*exp(-t_sat_/T1), to determine both k, the pseudo first-order forward reaction PCr→ATP rate constant (k_PCr_→_ATP_), and T1, the apparent T1 of PCr^124^.

### Quantitative reverse transcription polymerase chain reaction (RT-qPCR)

Brain tissues were harvested following decapitation under deep anaesthesia and immediately frozen onto dry ice. Total RNA was then extracted and purified using RNeasy Mini Kit (Qiagen, Venolo, Netherland) according to the vendor’s manual. Total RNA was then quantified and purity assessed using UV spectrophotometry. Reverse transcription was then performed on the samples using Taqman RT kit using Multiscribe Reverse Transcriptase (Thermofisher) and random hexamers. RT-qPCR was performed on the resulting cDNA samples at 0.16 ng/uL on 96-well plate with SYBR Green PCR Master Mix (Applied Biosystems). The reaction included a 2 min step at 50°C, a 10min step at 95°C, followed by 45 cycles of 15 s at 95°C – 1 min at 60°C – 30 s at 72°C. Relative gene expression was assessed with the comparative ΔΔCt method and normalized to both β-actine and β2 microglobulin (β-2m) as housekeeping genes. The primer list is described in Supplementary Table S2.

### Transcriptomics

Single-nucleus RNA sequencing (snRNA-seq) was performed on flash-frozen hippocampal tissue samples, with each sample comprising pooled material from five or six subjects. Hippocampal tissue was harvested following decapitation under terminal anaesthesia and flash-frozen on dry ice. Nuclei were isolated using the LeviPrep Nuclei Kit (LevitasBio). The nuclei were processed using the 10x Genomics Chromium X platform and the Next GEM 3ʹ assay according to the manufacturer’s protocol. Libraries were generated following 10x recommendations and sequenced on an Illumina platform by Novogene. Downstream data processing, including alignment to the reference genome, UMI counting, and gene expression quantification, was carried out using Cell Ranger software (10x Genomics).

The resulting snRNA-seq data were processed using *Scanpy* (v1.11.3)^125^. Raw gene–barcode matrices from each sample were imported using read_10x_mtx, and sample identity was stored in the AnnData metadata. Doublet detection was performed on each dataset using Scrublet^126^. Datasets were concatenated using adata.concatenate with genotype used as the batch identifier. Cells expressing fewer than 300 genes and genes detected in fewer than three cells were filtered out. Mitochondrial, ribosomal, and haemoglobin genes were annotated, and quality-control metrics were computed using sc.pp.calculate_qc_metrics. Cells with >6000 detected genes, >5% mitochondrial content, or >2% ribosomal content were excluded. Counts were normalized to 10,000 UMIs per cell and log-transformed using sc.pp.log1p. Highly variable genes were identified across batches (n = 12,000) using the method of Satija et al. (2015)^127^ as implemented in Scanpy. Total UMI counts and mitochondrial percentages were regressed out, and data were scaled to unit variance. Dimensionality reduction was performed via principal component analysis (PCA) using the *arpack* solver. Batch correction across genotypes was carried out using Harmony^128^, and Harmony-corrected principal components were used to compute the neighbourhood graph. UMAP embeddings were generated for visualization^129^, and unsupervised clustering was performed using the Leiden algorithm^130^ at a resolution of 0.6. Clusters were annotated manually based on canonical marker genes, and final cell-type labels were stored in the AnnData object. Cell-specific markers included *Prox1* for dentate gyrus neurons, *Grik4*, *Elavl2* and *Ccbe1* for Cornu Ammonis subfield 3 (CA) neurons, *Mpped1*, *Wfs1* and *Sv2b* for Cornu Ammonis subfield 1 and 2 (CA1/2), *Gad1* and *Gad2* for GABAergic neurons, *Gfap*, *Glul* and *Slc1a3* for astrocytes, *Sox10, Mbp* and *Plp1* for oligodendrocytes, *Hexb*, *Dock8* and *Cx3cr1* for microglia, *Pdgfra*, *Cspg4* and *Vcan* for Oligodendrocyte precursor cells (OPC), *Lef1*, *Flt1* and *Abcb1a* for endothelial cells, *Pdgfrb* for pericytes, *Reln* and *Trp73* for Cajal-Reztsius cells, *Foxj1* for ependymal cells. Single-cell gene set scores were computed using curated Gene Ontology (GO) Biological Process and Molecular Function gene sets related to insulin signalling (12 gene sets) or relevant to brain function (1,062 gene sets). Genes expressed in fewer than five cells were excluded to reduce noise. For each GO term, only genes present in the dataset and represented by at least four genes were retained. Gene set scores were calculated at the single-cell level using the Scanpy *sc.tl.score_genes* method, which computes the average expression of genes within a given gene set for each cell, relative to a reference gene pool. Within each cell type, mean gene set scores were computed separately for control and knockout cells and subtracted to generate a GO term × cell-type matrix summarizing differential pathway activity for downstream visualization and clustering. Hierarchical clustering was applied to the GO term × cell type differential score matrix using Ward’s minimum variance method with Euclidean distance, as implemented in seaborn’s clustermap function. Clustering was performed on GO terms (rows) to identify groups of functionally related pathways exhibiting similar patterns of regulation across cell types.

### Corticosterone administration

Mice were administered a subcutaneous dose (20 mg/kg) of corticosterone (CORT; VETRANAL^®^, CAS No. 50-22-6, Sigma-Aldrich) daily for 20 consecutive days. CORT was prepared as an emulsion (4mg/ml) in sesame oil containing 2% DMSO (CAS No. 67-68-5, Sigma-Aldrich). The vehicle group was administered the same solution without CORT. Animal’s body weight and welfare was monitored daily using score sheets. Cardboard sizzle nests were used for cage enrichment, and mice were handled using a cardboard tunnel to minimize stress associated with repeated injections^131^.

### Statistics

Statistical analyses were performed using GraphPad Prism 10 (GraphPad Software, San Diego, CA, USA), unless otherwise noted. Sample sizes were determined based on prior studies^19,132^. Two-group comparisons were analysed with two-tailed Student’s *t*-tests for normally distributed data, or two-tailed Mann–Whitney U tests when data were non-normal. The primary outcome pre-registered hypothesis (Glu/GABA ratio) in the humans study was tested with a one-tailed Student’s *t*-test. Multi-factor comparisons were performed using two-way ANOVA, with Bonferroni post-hoc tests applied for multiple comparisons when appropriate. Correlations were assessed using Pearson or Spearman correlation coefficients, as dictated by normality. Linear regression and principal component analysis (PCA) were performed within Prism. A significance threshold of *P* < 0.05 was applied. Fitting of creatine kinase (CK) rate constants (*kf*) was performed in MATLAB (R2020a) using a Levenberg–Marquardt algorithm. Each data point represents an individual animal or participant. All values are reported as mean ± s.d. unless otherwise specified. Detailed statistical tests are described in the figure legends or relevant methods sections.

## Supporting information

Supplementary Figues and Tables

## Acknowledgements

This study was supported financially by the Swiss National Science Foundation (P500PM_203208 to AC) and the John Fell Oxford University Press Research Fund (0016166 to AC). The Wellcome Centre for Integrative Neuroimaging (WIN, now OxCIN) was supported by core funding from the Wellcome Trust (203139/Z/16/Z and 203139/A/16/Z). This work was supported by the NIHR Oxford Health Biomedical Research Centre (NIHR203316). The views expressed are those of the author(s) and not necessarily those of the NIHR or the Department of Health and Social Care. We also thank the UK BBSRC (grant number BB/W019582/1) for support. WTC is funded by Wellcome [225924/Z/22/Z]. CJS holds a Wellcome Trust Senior Research Fellowship [224430/Z/21/Z]. The human MRS package was developed by Edward J. Auerbach and Małgorzata Marjańska and provided by the University of Minnesota under a C2P agreement.

We thank all the participants, as well as Jon Campbell, Michael Sanders, David Parker and Samira Abdelhamid for expert radiographic assistance and care of the participants during MRI scanning. We thank Stuart Pierson (NDCN, Oxford), Ma’ayan Semo (NDCN, Oxford), Katie Hewitt (Experimental Psychology department, Oxford) for support with preclinical experiments. We thank Piergiorgio Salvan (NDCN, Oxford) for support with rodent fMRI data processing.

This research was funded in whole, or in part, by the Wellcome Trust [Grant number 203139/Z/16/Z, 203139/A/16/Z, 224430/Z/21/Z, and 225924/Z/22/Z]. For the purpose of open access, the author has applied a CC BY public copyright licence to any Author Accepted Manuscript version arising from this submission.

## Authors’ contributions

AC, BRG, PC and JPL designed the study. AC, BRG, MT, MG, SZ, WTC, DB and SS optimized the protocol. AC, MG acquired the data. AC, AL, SZ, GD analysed the data. All authors interpreted the data. AC drafted the manuscript. All the authors assisted in revising the manuscript and approved the final version.

## Competing interests

The authors declare no conflicts of interest with respect to the research, authorship, and/or publication of this article.

## Supplementary information

Supplementary Information is available for this paper.

## Data availability statement

Source data underlying the figures will be made public upon publication. Single-nucleus RNA sequencing (snRNA-seq) data have been deposited in the European Nucleotide Archive (ENA) and BioStudies and will be made public upon publication. Additional data supporting the findings of this study are available from the corresponding author upon reasonable request.

## Notes

### Competing Interest Statement

The authors have declared no competing interest.

