## Supplementary Figues and Tables for "A hippocampal neuroimaging signature of neurovascular insulin signalling links metabolism to mood"

### Supplementary Figures

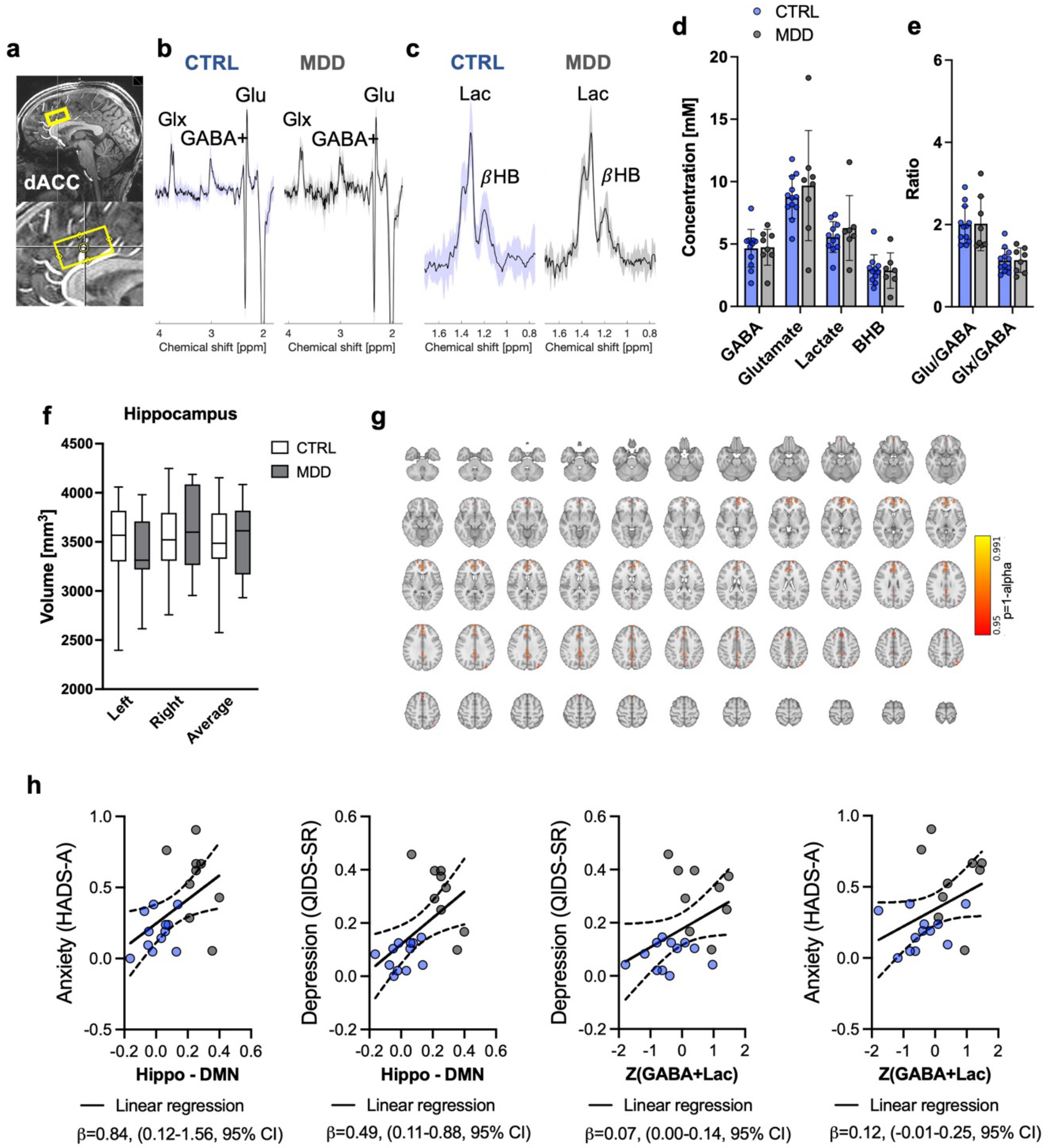

Supplementary Figure S1: *ACC function and metabolism are not associated with glycaemic variation and mood symptoms in depression*

***a.-c.*** *Dorsal anterior cingulate cortex (dACC) (**a**) voxel placement and average spectra (±s.d) for (**b**) GABA-edited and (**c**) lactate-edited $^1$H-MRS in CTRL (N=12, blue) and MDD (N=9, grey) groups.*

***d.,e.*** *Quantification of metabolites in dACC from $^1$H-MRS spectra.*

***f.*** *Hippocampal volume in CTRL and MDD subjects.*

***g.*** *Lightbox view of anterior hippocampal resting-state functional connectivity highlighting differences across groups (MDD>CTRL).*

***h.*** *Selected mood relationships to brain and metabolic metrics. Linear regressions with 95% CI.*

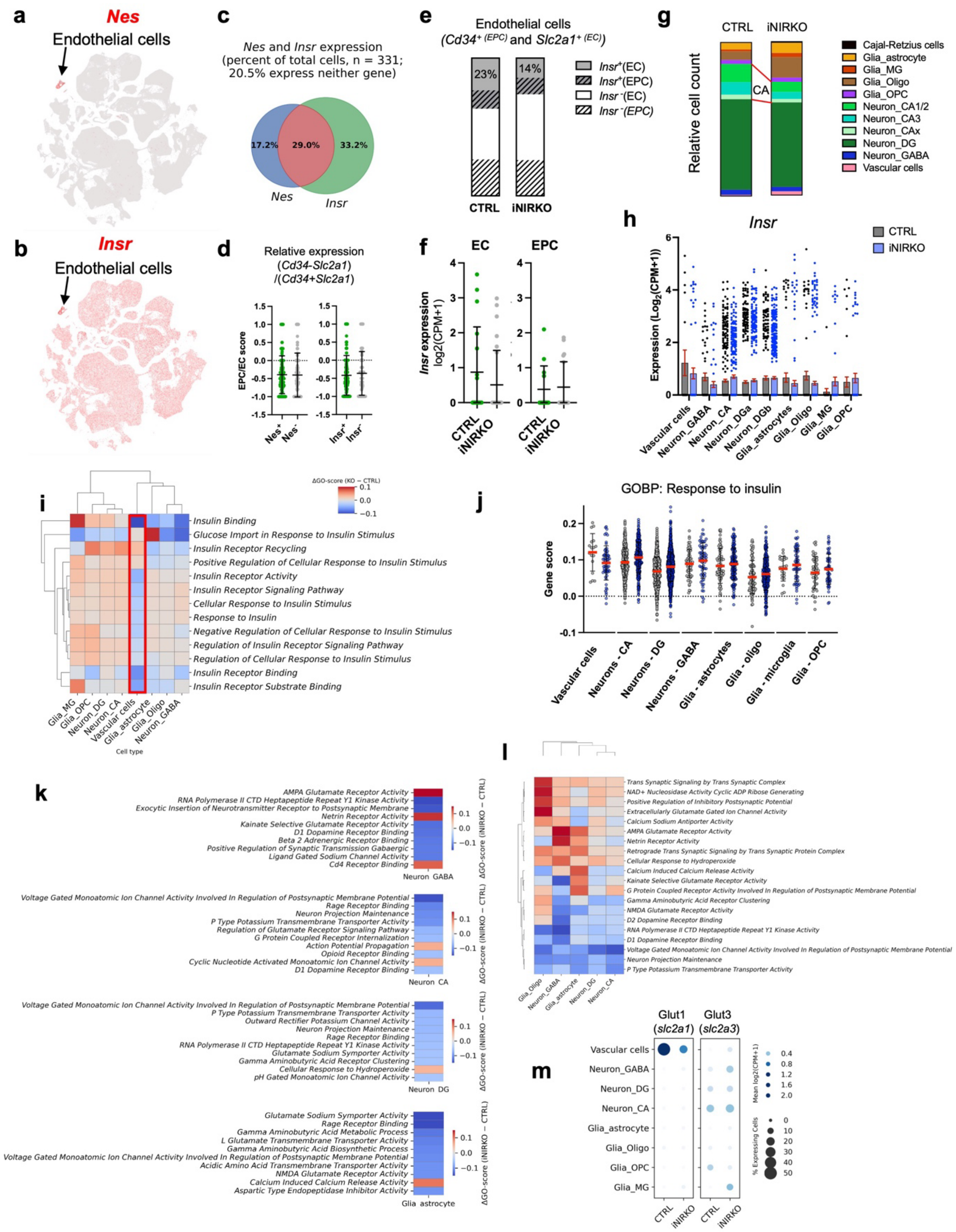
a
Nes
Endothelial cells
b
Insr
Endothelial cells
c
Nes and Insr expression
(percent of total cells, n = 331;
20.5% express neither gene)
17.2%
29.0%
33.2%
Nes
Insr
d
Relative expression
(Cd34-Slc2a1)
/(Cd34+Slc2a1)
EPC/EC score
Nes+
Nes-
Insr+
Insr-
e
Endothelial cells
(Cd34+ (EPC) and Slc2a1+ (EC))
23%
14%
Insr+(EC)
Insr+(EPC)
Insr-(EC)
Insr-(EPC)
CTRL
iNIRKO
f
EC
EPC
Insr expression
log2(CPM+1)
CTRL
iNIRKO
g
CTRL
iNIRKO
Relative cell count
CA
Cajal-Retzius cells
Glia_astrocyte
Glia_MG
Glia_Oligo
Glia_OPC
Neuron_CA1/2
Neuron_CA3
Neuron_CAx
Neuron_DG
Neuron_GABA
Vascular cells
h
Insr
CTRL
iNIRKO
Expression (Log2(CPM+1))
Vascular cells
Neuron_GABA
Neuron_CA
Neuron_DGa
Neuron_DGb
Glia_astrocytes
Glia_Oligo
Glia_MG
Glia_OPC
i
ΔGO-score (KO − CTRL)
Insulin Binding
Glucose Import in Response to Insulin Stimulus
Insulin Receptor Recycling
Positive Regulation of Cellular Response to Insulin Stimulus
Insulin Receptor Activity
Insulin Receptor Signaling Pathway
Cellular Response to Insulin Stimulus
Response to Insulin
Negative Regulation of Cellular Response to Insulin Stimulus
Regulation of Insulin Receptor Signaling Pathway
Regulation of Cellular Response to Insulin Stimulus
Insulin Receptor Binding
Insulin Receptor Substrate Binding
Glia_MG
Glia_OPC
Neuron_DG
Neuron_CA
Vascular cells
Glia_astrocyte
Glia_Oligo
Neuron_GABA
Cell type
j
GOBP: Response to insulin
Gene score
Vascular cells
Neurons - CA
Neurons - DG
Neurons - GABA
Glia - astrocytes
Glia - oligo
Glia - microglia
Glia - OPC
k
AMPA Glutamate Receptor Activity
RNA Polymerase II CTD Heptapeptide Repeat Y1 Kinase Activity
Exocytic Insertion of Neurotransmitter Receptor to Postsynaptic Membrane
Netrin Receptor Activity
Kainate Selective Glutamate Receptor Activity
D1 Dopamine Receptor Binding
Beta 2 Adrenergic Receptor Binding
Positive Regulation of Synaptic Transmission Gabaergic
Ligand Gated Sodium Channel Activity
Cd4 Receptor Binding
Neuron GABA
Voltage Gated Monoatomic Ion Channel Activity Involved In Regulation of Postsynaptic Membrane Potential
Rage Receptor Binding
Neuron Projection Maintenance
P Type Potassium Transmembrane Transporter Activity
Regulation of Glutamate Receptor Signaling Pathway
G Protein Coupled Receptor Internalization
Action Potential Propagation
Opioid Receptor Binding
Cyclic Nucleotide Activated Monoatomic Ion Channel Activity
D1 Dopamine Receptor Binding
Neuron CA
Voltage Gated Monoatomic Ion Channel Activity Involved In Regulation of Postsynaptic Membrane Potential
P Type Potassium Transmembrane Transporter Activity
Outward Rectifier Potassium Channel Activity
Neuron Projection Maintenance
Rage Receptor Binding
RNA Polymerase II CTD Heptapeptide Repeat Y1 Kinase Activity
Glutamate Sodium Symporter Activity
Gamma Aminobutyric Acid Receptor Clustering
Cellular Response to Hydroperoxide
pH Gated Monoatomic Ion Channel Activity
Neuron DG
Glutamate Sodium Symporter Activity
Rage Receptor Binding
Gamma Aminobutyric Acid Metabolic Process
L Glutamate Transmembrane Transporter Activity
Gamma Aminobutyric Acid Biosynthetic Process
Voltage Gated Monoatomic Ion Channel Activity Involved In Regulation of Postsynaptic Membrane Potential
Acidic Amino Acid Transmembrane Transporter Activity
NMDA Glutamate Receptor Activity
Calcium Induced Calcium Release Activity
Aspartic Type Endopeptidase Inhibitor Activity
Glia astrocyte
ΔGO-score (iNIRKO − CTRL)
l
Trans Synaptic Signaling by Trans Synaptic Complex
NAD+ Nucleosidase Activity Cyclic ADP Ribose Generating
Positive Regulation of Inhibitory Postsynaptic Potential
Extracellularly Glutamate Gated Ion Channel Activity
Calcium Sodium Antiporter Activity
AMPA Glutamate Receptor Activity
Netrin Receptor Activity
Retrograde Trans Synaptic Signaling by Trans Synaptic Protein Complex
Cellular Response to Hydroperoxide
Calcium Induced Calcium Release Activity
Kainate Selective Glutamate Receptor Activity
G Protein Coupled Receptor Activity Involved In Regulation of Postsynaptic Membrane Potential
Gamma Aminobutyric Acid Receptor Clustering
NMDA Glutamate Receptor Activity
D2 Dopamine Receptor Binding
RNA Polymerase II CTD Heptapeptide Repeat Y1 Kinase Activity
D1 Dopamine Receptor Binding
Voltage Gated Monoatomic Ion Channel Activity Involved In Regulation of Postsynaptic Membrane Potential
Neuron Projection Maintenance
P Type Potassium Transmembrane Transporter Activity
Glia_Oligo
Neuron_GABA
Glia_astrocyte
Neuron_DG
Neuron_CA
m
Glut1
(slc2a1)
Glut3
(slc2a3)
Vascular cells
Neuron_GABA
Neuron_DG
Neuron_CA
Glia_astrocyte
Glia_Oligo
Glia_OPC
Glia_MG
CTRL
iNIRKO
Mean log2(CPM+1)
% Expressing Cells

Supplementary Figure S2: *Non-parenchymal insulin receptor loss in the hippocampus models local brain insulin resistance and enhances neuronal insulin-responsive gene expression*

***a.-d.*** *Nestin is highly expressed in mature endothelial cells (EC). Uniform manifold approximation and projection (UMAP) of mouse whole cortex and hippocampus transcriptomics with expression (red) of (**a**) nestin (Nes) and (**b**) insulin receptor (Insr). (**c**) Overlap of Nes and Insr expression in endothelial cells. (**d**) Distribution of Nes and Insr expression is comparable across mature endothelial cell (EC; Slc2a1 positive) and endothelial progenitor cell (EPC; Cd34 positive). Data from Allen brain map (https://celltypes.brain-map.org/rnaseq/mouse_ctx-hpf_10x) and shown as relative expression of EC (Glut1/Slc2a1) and EPC (Cd34) gene markers (**c,d**).*

***e.-l.*** *Cell-specific changes in insulin signalling assessed with single nucleus RNAseq in dorsal hippocampus of CTRL (5 pooled samples) and iNIRKO (6 pooled samples) mice 6 weeks post tamoxifen induction. (**e**) Proportion of Insr expressing cells among Slc2a1 (EC) and Cd34 (EPC) positive cells. (**f**) Average expression of Insr among EC and EPC across genotypes. (**g**) Relative count for each cell type following UMAP and clustering. (**h**) Normalised expression of Insr. Data are mean ± s.e.m. (**i**) Gene score differences (iNIRKO - CTRL) across insulin-related gene ontology (GO) terms for all cell types. (**j**) Gene scores of GOBP: Response to insulin (GO:0032868) for all cell types. (**k**) Top 10 gene score differences (iNIRKO - CTRL) for neurons and astrocytes. (**l**) Top 20 (2%) Gene score differences (iNIRKO - CTRL) across cell types. (**m**) Dotplot of main glucose transporters Glut1 and Glut3 (Glut2 and Glut4 not detected) gene expression across cell types in the dorsal hippocampus.*

*Subfields of the hippocampus: CA, Cornu ammonis; DG, Dentate gyrus. Data are mean ± s.d. unless stated otherwise*

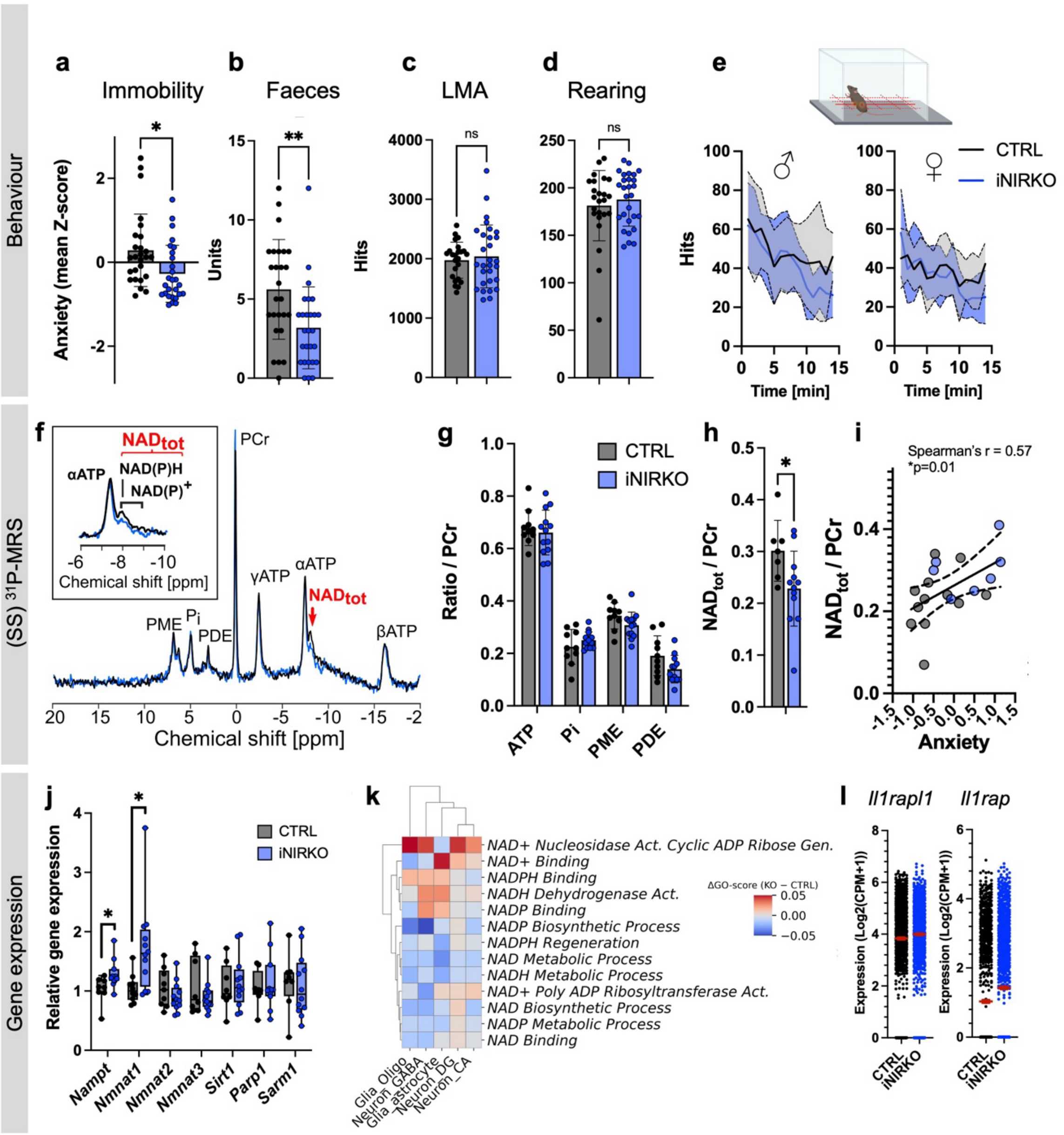
Behaviour
a
Immobility
Anxiety (mean Z-score)
b
Faeces
Units
c
LMA
Hits
ns
d
Rearing
Hits
ns
e
Hits
Time [min]
CTRL
iNIRKO
(SS) 31P-MRS
f
NADtot
αATP
NAD(P)H
NAD(P)+
Chemical shift [ppm]
PCr
γATP
αATP
NADtot
PME
Pi
PDE
βATP
Chemical shift [ppm]
g
Ratio / PCr
CTRL
iNIRKO
ATP
Pi
PME
PDE
h
NADtot / PCr
i
Spearman's r = 0.57
*p=0.01
NADtot / PCr
Anxiety
Gene expression
j
Relative gene expression
CTRL
iNIRKO
Nampt
Nmnat1
Nmnat2
Nmnat3
Sirt1
Parp1
Sarm1
k
NAD+ Nucleosidase Act. Cyclic ADP Ribose Gen.
NAD+ Binding
NADPH Binding
NADH Dehydrogenase Act.
NADP Binding
NADP Biosynthetic Process
NADPH Regeneration
NAD Metabolic Process
NADH Metabolic Process
NAD+ Poly ADP Ribosyltransferase Act.
NAD Biosynthetic Process
NADP Metabolic Process
NAD Binding
ΔGO-score (KO − CTRL)
Glia_Oligo
Neuron_GABA
Glia_astrocyte
Neuron_DG
Neuron_CA
l
Il1rapl1
Il1rap
Expression (Log2(CPM+1))
CTRL
iNIRKO

Supplementary Figure S3: *Non-parenchymal insulin resistance increases metabolic activity and function in the hippocampus and has anxiolytic-like effects*

***a.*** *Average immobility measured in EPM and AOF tests. Student's t-test, *p<0.05,*

***b.*** *Average defecation during in EPM, LDB and AOF tests. Mann-Whitney test, **p<0.005;*

***c.,d.,e,*** *Locomotor activity measured for 60min in the monitoring cage with average number of photobeam hits (**c**) in the x-y (LMS) or (**d**) or z dimensions (rearing) of the cage. (**e**) Genotype and sex-specific locomotor activity during the first 15min of the test in the x-y dimension.*

***f.*** *Steady-state (SS) $^{31}$P-MRS average spectra for iNIRKO (blue) and CTRL (black) mice with a zoom on the $NAD_{tot}$ peaks.*

***g.h.*** *(**g**) Energy (ATP, Pi), membrane (PME, PDE) and (**h**) redox (NAD) metabolite quantifications from (SS) $^{31}$P-MRS spectra. Student's t-test, *p<0.05,*

***i.*** *Relationship of $NAD_{tot}$ with anxiety score. Correlation (*p <0.05) and simple linear regression fit ($\beta$=0.055) shown with 95% CI.*

***f.*** *Relative NAD(H)-related gene expression in the dorsal hippocampus of iNIRKO and CTRL mice measured with qPCR. iNIRKO, n=12; CTRL, n=8. Student's t-test, *p<0.05.*

***k.*** *Gene score differences (iNIRKO - CTRL) for NAD(H)-related processes across cell types.*

***l.*** *Average expression of top NAD(H)-related genes in all cell types. Data are mean $\pm$ s.e.m*

*All data are mean $\pm$ s.d. unless stated otherwise*

*LMA, locomotor activity; EPM, elevated plus maze; AOF, Aversive Open-field test; LDB, light-dark box test; NAD, Nicotinamide adenine dinucleotide; ATP, Adenosine triphosphate; PCr, phosphocreatine; Pi, inorganic phosphate; PME, phosphomonoesters; PDE, phosphodiesters.*

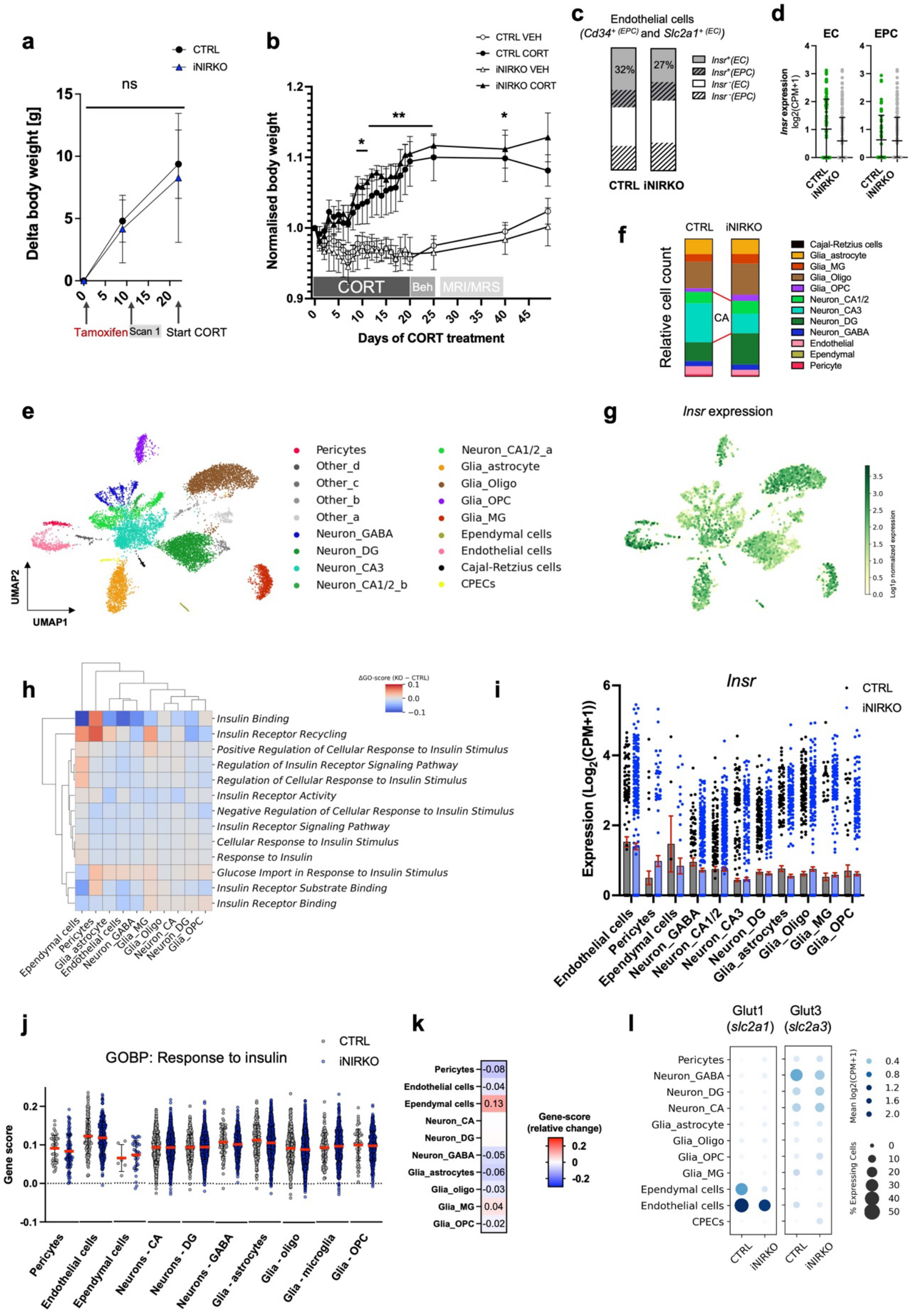
a
CTRL
iNIRKO
ns
Delta body weight [g]
Tamoxifen
Scan 1
Start CORT
b
CTRL VEH
CTRL CORT
iNIRKO VEH
iNIRKO CORT
Normalised body weight
CORT
Beh
MRI/MRS
Days of CORT treatment
c
Endothelial cells
(Cd34+ (EPC) and Slc2a1+ (EC))
32%
27%
Insr+(EC)
Insr+(EPC)
Insr−(EC)
Insr−(EPC)
CTRL iNIRKO
d
EC
EPC
Insr expression log2(CPM+1)
CTRL
iNIRKO
e
Pericytes
Other_d
Other_c
Other_b
Other_a
Neuron_GABA
Neuron_DG
Neuron_CA3
Neuron_CA1/2_b
Neuron_CA1/2_a
Glia_astrocyte
Glia_Oligo
Glia_OPC
Glia_MG
Ependymal cells
Endothelial cells
Cajal-Retzius cells
CPECs
UMAP2
UMAP1
f
CTRL
iNIRKO
Relative cell count
CA
Cajal-Retzius cells
Glia_astrocyte
Glia_MG
Glia_Oligo
Glia_OPC
Neuron_CA1/2
Neuron_CA3
Neuron_DG
Neuron_GABA
Endothelial
Ependymal
Pericyte
g
Insr expression
Log1p normalized expression
h
ΔGO-score (KO − CTRL)
Insulin Binding
Insulin Receptor Recycling
Positive Regulation of Cellular Response to Insulin Stimulus
Regulation of Insulin Receptor Signaling Pathway
Regulation of Cellular Response to Insulin Stimulus
Insulin Receptor Activity
Negative Regulation of Cellular Response to Insulin Stimulus
Insulin Receptor Signaling Pathway
Cellular Response to Insulin Stimulus
Response to Insulin
Glucose Import in Response to Insulin Stimulus
Insulin Receptor Substrate Binding
Insulin Receptor Binding
Ependymal cells
Pericytes
Glia_astrocyte
Endothelial cells
Neuron_GABA
Glia_MG
Glia_Oligo
Neuron_CA
Neuron_DG
Glia_OPC
i
Insr
CTRL
iNIRKO
Expression (Log2(CPM+1))
Endothelial cells
Pericytes
Ependymal cells
Neuron_GABA
Neuron_CA1/2
Neuron_CA3
Neuron_DG
Glia_astrocytes
Glia_Oligo
Glia_MG
Glia_OPC
j
GOBP: Response to insulin
CTRL
iNIRKO
Gene score
Pericytes
Endothelial cells
Ependymal cells
Neurons - CA
Neurons - DG
Neurons - GABA
Glia - astrocytes
Glia - oligo
Glia - microglia
Glia - OPC
k
Pericytes -0.08
Endothelial cells -0.04
Ependymal cells 0.13
Neuron_CA
Neuron_DG
Neuron_GABA -0.05
Glia_astrocytes -0.06
Glia_oligo -0.03
Glia_MG 0.04
Glia_OPC -0.02
Gene-score (relative change)
l
Glut1 (slc2a1)
Glut3 (slc2a3)
Pericytes
Neuron_GABA
Neuron_DG
Neuron_CA
Glia_astrocyte
Glia_Oligo
Glia_OPC
Glia_MG
Ependymal cells
Endothelial cells
CPECs
CTRL
iNIRKO
Mean log2(CPM+1)
% Expressing Cells

**Supplementary Figure S4:** *Hippocampal non-parenchymal insulin resistance stabilizes long-term metabolic and behavioural trajectories*

***a.,b.*** *Evolution of body weight following tamoxifen induction before (**a**) and after (**b**) the start of corticosterone administration. Repeated measure two-way ANOVA, (**a**), ns, not significant; (**b**) Treatment effect: $F_{1,25} = 22.95$, $P < 0.0001$, Bonferroni's posthoc test *$p<0.05$; **$p<0.005$; Data are mean $\pm$ s.e.m.*

***c-j**. Cell-specific changes in insulin signalling assessed with single nucleus RNAseq of whole hippocampus of CTRL (6 pooled samples) and iNIRKO (6 pooled samples) mice 23 weeks post tamoxifen. (**c**) Proportion of Insr expressing cells among Slc2a1 (EC) and Cd34 (EPC) positive cells. (**d**) Average expression of Insr among EC and EPC across genotypes. (**e**) Uniform manifold approximation and projection (UMAP) with (**f**) relative count for each cell type and (**g**) Insr expression for iNIRKO (n=10,296 cells, 6 pooled samples) and CTRL (n=2,515 cells, 6 pooled samples) mice (VEH groups) combined. (**h**) Gene score differences (iNIRKO - CTRL) across insulin-related gene ontology (GO) terms for all cell types. (**i**) Normalised expression of Insr. Data are mean $\pm$ s.e.m. Absolute (**j**) and relative (**k**) change change (iNIRKO-CTRL) in gene score for the reponse to insulin GO term (GO:0032868). (**l**) Dotplot of Glu1 and Glu3 gene expression across cell types.*

*CORT, corticosterone; VEH, vehicle; EC, endothelial cell; EPC, endothelial progenitor cell.*

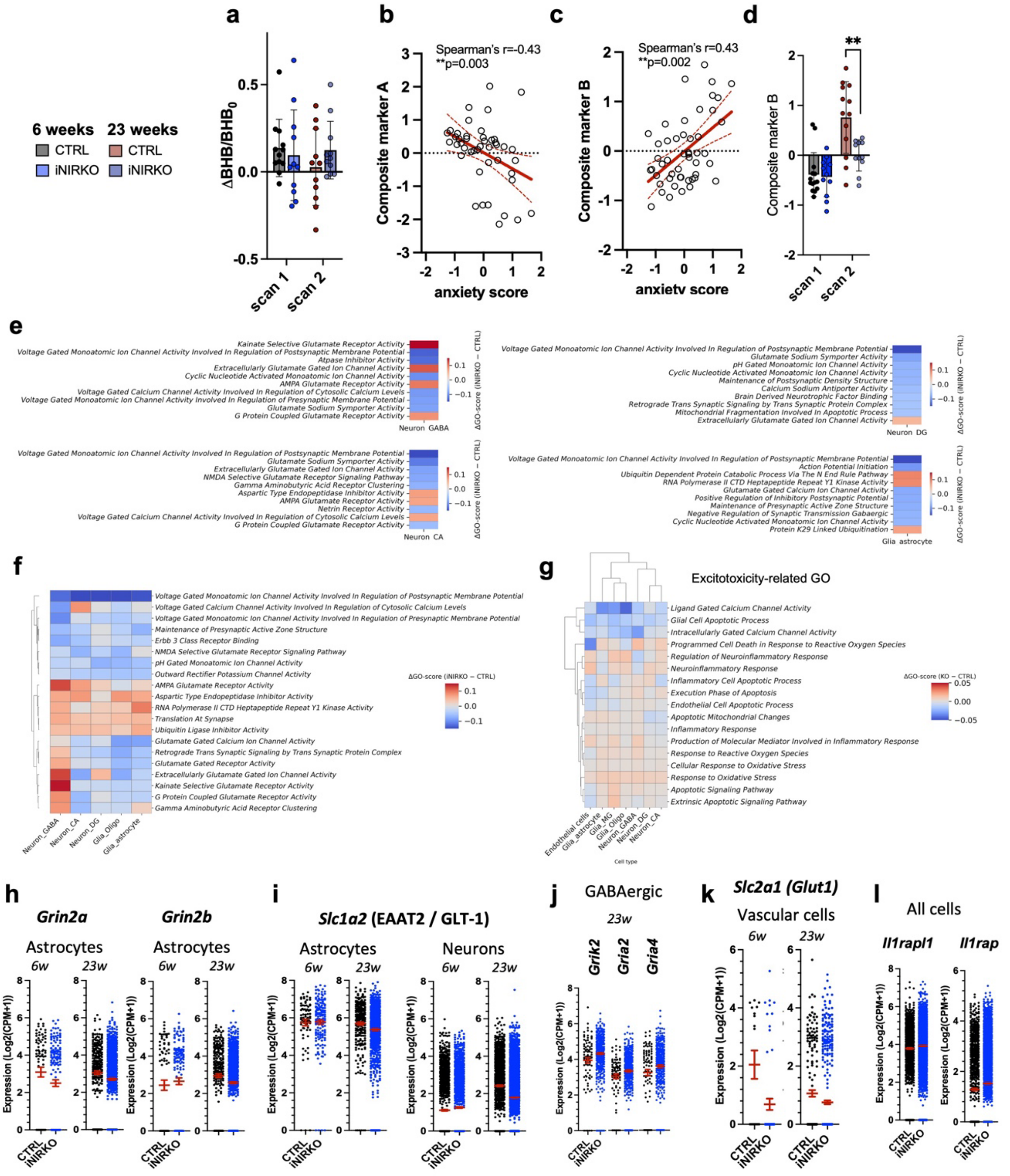
a
b
c
d
6 weeks
23 weeks
CTRL
iNIRKO
CTRL
iNIRKO
ΔBHB/BHB0
scan 1
scan 2
Spearman's r=-0.43
**p=0.003
Composite marker A
anxiety score
Spearman's r=0.43
**p=0.002
Composite marker B
anxiety score
Composite marker B
**
e
Kainate Selective Glutamate Receptor Activity
Voltage Gated Monoatomic Ion Channel Activity Involved In Regulation of Postsynaptic Membrane Potential
Atpase Inhibitor Activity
Extracellularly Glutamate Gated Ion Channel Activity
Cyclic Nucleotide Activated Monoatomic Ion Channel Activity
AMPA Glutamate Receptor Activity
Voltage Gated Calcium Channel Activity Involved In Regulation of Cytosolic Calcium Levels
Voltage Gated Monoatomic Ion Channel Activity Involved In Regulation of Presynaptic Membrane Potential
Glutamate Sodium Symporter Activity
G Protein Coupled Glutamate Receptor Activity
Neuron GABA
ΔGO-score (iNIRKO − CTRL)
Voltage Gated Monoatomic Ion Channel Activity Involved In Regulation of Postsynaptic Membrane Potential
Glutamate Sodium Symporter Activity
pH Gated Monoatomic Ion Channel Activity
Cyclic Nucleotide Activated Monoatomic Ion Channel Activity
Maintenance of Postsynaptic Density Structure
Calcium Sodium Antiporter Activity
Brain Derived Neurotrophic Factor Binding
Retrograde Trans Synaptic Signaling by Trans Synaptic Protein Complex
Mitochondrial Fragmentation Involved In Apoptotic Process
Extracellularly Glutamate Gated Ion Channel Activity
Neuron DG
Voltage Gated Monoatomic Ion Channel Activity Involved In Regulation of Postsynaptic Membrane Potential
Glutamate Sodium Symporter Activity
Extracellularly Glutamate Gated Ion Channel Activity
NMDA Selective Glutamate Receptor Signaling Pathway
Gamma Aminobutyric Acid Receptor Clustering
Aspartic Type Endopeptidase Inhibitor Activity
AMPA Glutamate Receptor Activity
Netrin Receptor Activity
Voltage Gated Calcium Channel Activity Involved In Regulation of Cytosolic Calcium Levels
G Protein Coupled Glutamate Receptor Activity
Neuron CA
Voltage Gated Monoatomic Ion Channel Activity Involved In Regulation of Postsynaptic Membrane Potential
Action Potential Initiation
Ubiquitin Dependent Protein Catabolic Process Via The N End Rule Pathway
RNA Polymerase II CTD Heptapeptide Repeat Y1 Kinase Activity
Glutamate Gated Calcium Ion Channel Activity
Positive Regulation of Inhibitory Postsynaptic Potential
Maintenance of Presynaptic Active Zone Structure
Negative Regulation of Synaptic Transmission Gabaergic
Cyclic Nucleotide Activated Monoatomic Ion Channel Activity
Protein K29 Linked Ubiquitination
Glia astrocyte
f
Voltage Gated Monoatomic Ion Channel Activity Involved In Regulation of Postsynaptic Membrane Potential
Voltage Gated Calcium Channel Activity Involved In Regulation of Cytosolic Calcium Levels
Voltage Gated Monoatomic Ion Channel Activity Involved In Regulation of Presynaptic Membrane Potential
Maintenance of Presynaptic Active Zone Structure
Erbb 3 Class Receptor Binding
NMDA Selective Glutamate Receptor Signaling Pathway
pH Gated Monoatomic Ion Channel Activity
Outward Rectifier Potassium Channel Activity
AMPA Glutamate Receptor Activity
Aspartic Type Endopeptidase Inhibitor Activity
RNA Polymerase II CTD Heptapeptide Repeat Y1 Kinase Activity
Translation At Synapse
Ubiquitin Ligase Inhibitor Activity
Glutamate Gated Calcium Ion Channel Activity
Retrograde Trans Synaptic Signaling by Trans Synaptic Protein Complex
Glutamate Gated Receptor Activity
Extracellularly Glutamate Gated Ion Channel Activity
Kainate Selective Glutamate Receptor Activity
G Protein Coupled Glutamate Receptor Activity
Gamma Aminobutyric Acid Receptor Clustering
ΔGO-score (iNIRKO − CTRL)
Neuron_GABA
Neuron_CA
Neuron_DG
Glia_Oligo
Glia_astrocyte
g
Excitotoxicity-related GO
Ligand Gated Calcium Channel Activity
Glial Cell Apoptotic Process
Intracellularly Gated Calcium Channel Activity
Programmed Cell Death in Response to Reactive Oxygen Species
Regulation of Neuroinflammatory Response
Neuroinflammatory Response
Inflammatory Cell Apoptotic Process
Execution Phase of Apoptosis
Endothelial Cell Apoptotic Process
Apoptotic Mitochondrial Changes
Inflammatory Response
Production of Molecular Mediator Involved in Inflammatory Response
Response to Reactive Oxygen Species
Cellular Response to Oxidative Stress
Response to Oxidative Stress
Apoptotic Signaling Pathway
Extrinsic Apoptotic Signaling Pathway
ΔGO-score (KO − CTRL)
Endothelial cells
Glia_astrocyte
Glia_MG
Glia_Oligo
Neuron_GABA
Neuron_DG
Neuron_CA
Cell type
h
Grin2a
Grin2b
Astrocytes
6w
23w
Expression (Log2(CPM+1))
CTRL
iNIRKO
i
Slc1a2 (EAAT2 / GLT-1)
Astrocytes
Neurons
j
GABAergic
23w
Grik2
Gria2
Gria4
k
Slc2a1 (Glut1)
Vascular cells
l
All cells
Il1rapl1
Il1rap

Supplementary Figure S5: Hippocampal insulin resistance limits age-dependent glutamate-driven metabolic changes

***a.*** *Relative difference in total BHB concentrations before and after* $^{13}$*C-glucose infusion at 6 and 23 weeks post tamoxifen induction.*

***b.,c.*** *Relationship of composite markers A and B with anxiety. Composite marker A (**b**) was defined as the z-score of the labelling ratio* $^{13}$*C-GluH4/*$^{13}$*C-GABAH4 multiplied with net lactate production (Δlactate/lactate*$_0$*). Composite marker B (**c**) was defined as the average of individual z-scores including* $^{13}$*C-GluH4/GlxH2FE, (Δlactate/lactate*$_0$*), GABA and lactate concentrations. Data from 6 and 23 weeks timepoints. Correlation (**p <0.005) and simple linear regression fit (*$\beta_{marker-A}$*=-0.48,* $\beta_{marker-B}$*=0.48) shown with 95% CI.*

***d.*** *Group differences in composite metabolic marker B at 6 and 23 weeks post tamoxifen induction. Student's t-test, **p<0.005.*

***e.*** *Top 20 (2%) Gene score differences (iNIRKO - CTRL) across cell types at 23 weeks post tamoxifen.*

***f.*** *Gene scores of excitotoxicity-related GO terms across cell types at 23 weeks post tamoxifen.*

***g.-h,*** *Average expression of genes driving GO differences in (**h**) calcium channel activity, (**i**) glutamate sodium symporter activity, (**j**) kainite selective glutamate transporter activity, (**k**) vascular glucose uptake and (**l**) NAD(H)-related GO terms. Data are mean* $\pm$ *s.d.*

### Supplementary Tables

| session | Breathing rate [rpm] | | Rectal Temperature [°C] | |
|---|---|---|---|---|
| | *CTRL* | *iNIRKO* | *CTRL* | *iNIRKO* |
| SS $^{31}$P-MRS | 143±22 | 157±20 | 36.3 ± 0.7 | 36.0 ± 1.0 |
| ST $^{31}$P-MRS | 148 ± 23 | 155 ± 18 | 36.4 ± 0.9 | 36.5 ± 0.6 |
| POPE$^{13}$C-MRS/rs-fMRI 1 | 166 ± 15 | 179 ± 19 | 36.9 ± 0.2 | 36.9 ± 0.3 |
| POPE$^{13}$C-MRS 2 | 161 ± 19 | 164 ± 15 | 37.1 ± 0.5 | 36.9 ± 0.6 |

**Supplementary Table S1:** Mouse MRI physiological parameters

| Gene | Forward sequence (5'-3') | Reverse sequence (5'-3') |
|---|---|---|
| *Insr E3-4* | AAATGCAGGAACTCTCGGAAGCCT | ACCTTCGAGGATTTGGCAGACCTT |
| *Insr E21* | TTTGTCATGGATGGAGGCTA | CCTCATCTTGGGGTTGAACT |
| *Insr A* | TCCTGAAGGAGCTGGAGGAGT | CTTTCGGGATGGCCTGG |
| *Insr B* | idem | TTCGGGATGGCCTACTGTC |
| *β-Actine* | GCTTCTTTGCAGCTCCTTCGT | ATATCGTCATCCATGGCGAAC |
| *β-2m* | TGTCAGATATGTCCTTCAGCAAGG | TGCTTAACTCTGCAGGCGTATG |
| *Nampt* | TCGGTTCTGGTGGCGCTTTGCTAC | AAGTTCCCCGCTGGTGTCCTATGT |
| *Nmnat1* | GAAATTGCTGTGTGGGGCAG | CCACGATTTGCGTGATGTCC |
| *Nmnat2* | GATGTTCGAGAGAGCCAGGG | AAGGCCCTGTTTTCCGTAGG |
| *Nmnat3* | AAGACACCATCAGCCTCTGC | CCAAGCCGAACTTCTCCACT |
| *Sirt1* | AGTTCCAGCCGTCTCTGTGT | GATCCTTTGGATTCCTGCAA |
| *Parp1* | GGCAAGCACAGTGTCAAAGG | TGTCGTTGACACCAGATGGG |
| *Sarm1* | CGCTGCCCTGTACTGGAGG | CTTCAGGAGGCTGGCCAGCT |

**Supplementary Table S2:** List of Primers

| Variable | CTRL (N = 12) | MDD (N = 9) | Statistics |
|---|---|---|---|
| Age (years) | 34±14 (Mean±SD) | 38 ± 12 (Mean±SD) | U=41, p=0.37 |
| Sex | females: 6 (50%), males: 6 (50%) | females: 3 (33.3%), males: 6 (66.7%) | $\chi^2$ =0.58, p=0.44 |
| Cognition (MoCA) | 28.83 ± 1.40 (Mean±SD) | 28.78 ± 1.48 (Mean±SD) | U=53, p=0.99 |
| BMI | 24.55 ± 2.75 | 25.34 ± 3.81 | t=0.53, p=0.60 |
| Body fat (%) | 30.58 ± 8.98 | 28.94 ± 9.06 | t=0.41, p=0.68 |
| Skeletal muscle (%) | 30.71 ± 5.38 | 31.73 ± 5.44 | t=0.43, p=0.67 |
| Visceral fat | 6.4 ± 2.5 | 8.1 ± 3.6 | t=1.21, p=0.25 |
| Resting metabolism [kcal] | 1,551 ± 180 | 1,622 ± 316 | t=0.61, p=0.55 |

**Supplementary Table S3:** Participants demographics

*Group comparisons were done with Welch's t-test normally distributed variables with unequal variances and the Mann-Whitney U tests for non-parametric variables. Sex distribution was compared a $\chi^2$ test.*
